# Single-cell spatial atlas of high-grade serous ovarian cancer unveils MHC class II as a key driver of spatial tumor ecosystems and clinical outcomes

**DOI:** 10.1101/2024.11.29.626039

**Authors:** Fernando Perez-Villatoro, Lilian van Wagensveld, Aleksandra Shabanova, Ada Junquera, Ziqi Kang, Iga Niemiec, Matias M Falco, Ella Anttila, Julia Casado, Eric Marcus, Essi Kahelin, Foteini Chamchougia, Matilda Salko, Saundarya Shah, Salvatore Russo, Jacopo Chiaro, Mikaela Grönholm, Gabe S. Sonke, Koen K. Van de Vijver, Rutgerus FPM Kruitwagen, Maaike Avan der Aa, Anni Virtanen, Vincenzo Cerullo, Anna Vähärautio, Peter K. Sorger, Hugo M. Horlings, Anniina Färkkilä

## Abstract

The tumor microenvironment (TME) is a complex network of interactions between malignant and host cells, yet its orchestration in advanced high-grade serous ovarian carcinoma (HGSC) remains poorly understood. We present a comprehensive single-cell spatial atlas of 280 metastatic HGSCs, integrating high-dimensional imaging, genomics, and transcriptomics. Using 929 single-cell maps, we identify distinct spatial domains associated with phenotypically heterogeneous cellular compositions, and demonstrate that immune cell co-infiltration at the tumor-stroma interface significantly influences clinical outcomes. To uncover the key drivers of the tumor ecosystem, we developed CEFIIRA (Cell Feature Importance Identification by RAndom forest), which identified tumor cell-intrinsic MHC-II expression as a critical predictor of prolonged survival, independent of clinicomolecular profiles. Validation with external datasets confirmed that MHC-II-expressing cancer cells drive immune infiltration and orchestrate spatial tumor-immune interactions. Our atlas offers novel insights into immune surveillance mechanisms across HGSC clinicomolecular groups, paving the way for improved therapeutic strategies and patient stratification.

## INTRODUCTION

The tumor microenvironment (TME) is a complex ecosystem comprising cancer cells, immune cells, stromal cells, and non-cellular components and their interactions^1^. The continuous crosstalk between tumor cells and the surrounding microenvironment is fundamental to tumor initiation, phenotypic changes, cancer progression, and therapeutic resistance. Ovarian high-grade serous carcinoma (HGSC) is the most prevalent and aggressive subtype of ovarian cancer ^2^, characterized by high genomic instability ^3^ and significant tumor heterogeneity ^4^. This heterogeneity is evident in various aspects, such as histopathological architecture, genetic alterations, and TME cellular composition, leading to varying responses to chemotherapy ^5^. The genetic diversity of HGSC tumor cells significantly influences TME. For instance, tumors with a homologous recombination (HR) DNA repair deficiency, such as those with *BRCA1/2* mutations, have been suggested to be more immunogenic and exhibit higher lymphocyte infiltration ^6,7^. Emerging evidence also suggests that HR-deficient (HRD) tumors possess distinct immune escape mechanisms ^8^ and unique responses to combinations with immunotherapies ^9,10^. Moreover, we and others have shown that exposure to neoadjuvant chemotherapy significantly shapes the tumor ecosystem by enhancing the intricate interactions within the immune compartment ^11–13^ resulting in spatially confined T-cell exhaustion^14^. Therefore, it is crucial to decode the complex relationships between clinicomolecular features and the tumor ecosystem to improve patient stratification and immunotherapeutic treatment strategies for patients with HGSC.

We developed a high-resolution multi-omics single-cell spatial atlas, analyzing 15.1 million cells from advanced HGSC patient samples with comprehensive clinical and follow-up data. Leveraging detailed DNA methylation, gene copy number, and mutational profiling, we established clinicomolecular profiles that reveal critical tumor ecosystem dynamics. Using our image analysis paradigm, we resolved the TME at the single-cell level, identifying orchestrated multicellular domains that shape patient outcomes. We employed a custom machine learning tool, CEFIIRA, to integrate single-cell spatial, molecular, and survival data to uncover survival-linked features. We highlight the pivotal role of cancer cell-intrinsic MHC class II (MHCII) expression specifically at the tumor-stroma interface in driving immune landscapes. We show that MHCII-positive cancer cells generate immune hotspots marked by increased activation, impacting prognosis independently of clinicomolecular profiles. This atlas decodes the interplay between tumor and immune cells, shedding light on mechanisms influencing prognosis and paving the way for targeted interventions in HGSC.

## RESULTS

### Multi-omics single-cell spatial mapping of the tumor ecosystem of HGSC

To uncover the relationship between the TME ecosystem topography, clinical prognosis, and molecular profiles in HGSC we applied multi-omics analysis combined with high-plex imaging at a subcellular resolution on 265 advanced HGSCs (**Fig. 1a**). The tumor tissues were obtained from debulking surgeries, either from treatment-naive patients (Primary debulking surgery, PDS n=100) or after platinum-based neoadjuvant chemotherapy (Interval debulking surgery, IDS n=158). The therapy sequence (PDS/IDS) was unknown for seven patients. Detailed clinical data, including long-term clinical outcomes, were collected, with a median follow-up of 38.2 months (IQR: 22-67 months) (**Supplementary Table 1**). Most patients (87.5%) received carboplatin and paclitaxel as adjuvant chemotherapy, and the median progression-free survival was 16 months (IQR: 12-27 months).

**Figure 1.**
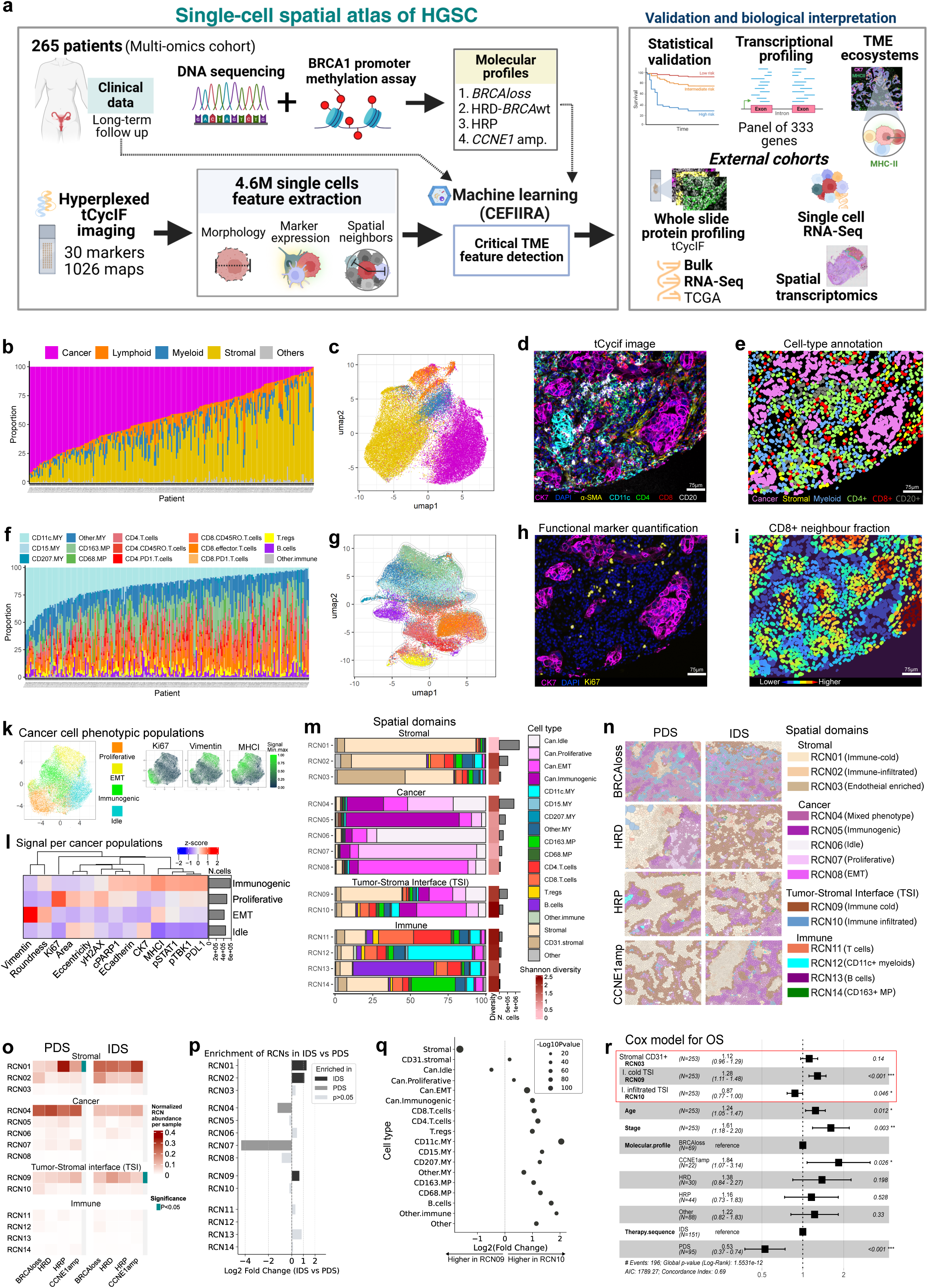
**a** Schematic of our study. We analyzed 265 HGSC patients using multi-omics tumor profiling. Each tumor was categorized in one of four common HGSC profiles: BRCAloss, homologous recombination deficient (HRD-BRCAwt), homologous recombination proficient (HRP) and Cyclin-E amplification (CCNE1 amp.), using methylation assay, targeted gene sequencing and shallow whole genome sequencing. The tumor microenvironment (TME) was assessed using tissue microarrays, comprising 1026 cores (spatial maps). High-dimensional imaging was employed to study the TME and extract single-cell spatial features. We developed a machine learning pipeline (CEFIIRA) to integrate the single-cell features and adjust for clinical covariates, aiming to predict features associated with survival and molecular profiles. Our results were validated using statistical methods, followed by further biological analysis for interpretation, both incorporating external cohorts. **b** A barplot displaying the composition of cancer cells, stromal cells, myeloid cells, lymphocytes, and unassigned cells per tumor. The barplot is sorted by cancer cell abundance. **c** UMAP projection of the analyzed cells, the colors correspond to their main cell type group assignment. **d.** Example of tCycIF image. **e** Cell type annotation of the image in panel d**. f** Barplot illustrating the composition of immune cells per tumor. The barplot is sorted according to the CD11c+ myeloid cell proportion. **g** UMAP projection of immune cells coloured by their annotation as in panel f. **h** tCycIF image displaying the functional marker Ki67; this image corresponds to the one in panel d. **i** Cells labeled according to their proportion of CD8.T cells neighbors, areas with higher CD8.T cell neighbors are shown in red, this image corresponds to the one in panel d. **k** UMAP projection of cancer cells, coloured by their cluster assignment (at the left) and by the expression of relevant proteins (smaller panels at the right). **l** Heatmap displaying the median expression of markers per cancer cluster. The listed markers were used for cancer-cell clustering. A hierarchical clustering of the markers based on their median expression per cluster is shown in the dendrograms. **m** Barplot displays the composition of cells in each recurrent cellular neighborhood (RCN). The adjacent small red boxes show their corresponding Shannon diversity. On the right, the gray barplots show the total number of cells assigned to each RCN. **n** RCNs across the molecular profiles and therapy sequences. **o** Heatmaps displaying the median proportion of cells assigned to the RCNs, per molecular profile. Individual heatmaps are presented for PDS and IDS samples. For each RNC’s, its differential abundance across the four molecular profiles was calculated using a two-tailed Kruskal-Wallis test. The significant differences (unadjusted P<0.05) are highlighted in teal boxes. **p** Barplots of the enrichment (log2 fold change) of RCNs in IDS samples compared to PDS. Enrichment was calculated using Wilcoxon two sided test. **q** Fold change in the proportion of cells in RCN09 compared to RCN10. The differential difference was calculated using Wilcoxon two sided test. **r** Forest plot of the Cox regression of overall survival in relation to the abundance of indicated RCNs and adjusted by the displayed variables.

We conducted a hierarchical multi-omics analysis to assign the tumors’ molecular profiles. Firstly, *BRCA1/2* germline or somatic mutation status was assessed using targeted high throughput sequencing, and *BRCA1* promoter hypermethylation was evaluated using a targeted DNA methylation assay. Tumors without BRCAloss (*BRCA1/2* somatic or germline mutation, or *BRCA1* promoter hypermethylation) were tested for Homologous Recombination Deficiency (HRD) and CCNE1-amplification using shallow whole genome sequencing and dedicated algorithms (see Methods). We categorized 174 tumors (**Supplementary Fig. 1a)** into four clinically and molecularly relevant groups (*BRCAloss, HRD, HRP, and CCNE1 amplification*) ^15^. The rest (n=91) of the tumors were left “unassigned” as they could not be reliably categorized into the predefined molecular profiles (see Methods). For gene expression profiling, we performed Nanostring analysis using a panel of 335 genes ^16^. To characterize the TME from this large cohort, a tissue microarray (TMA) was constructed by targeting two representative cores from the tumor and two from stromal-rich areas. The tumor tissues were also stained for H&E and CD8, CD20, CD103, CD68 using immunohistochemistry (IHC).

To precisely map the spatial TME at a single-cell resolution, we used hyperplexed imaging with tissue cyclic immunofluorescence (tCycIF) targeting 30 different proteins (**Supplementary Table 2**), generating 929 single-cell maps of HGSC (**Fig. 1a**). Image processing and analysis yielded 4,644,485 high-quality single cells, with an average of 20,000 cells for each tumor. The cell phenotyping was based on the multidimensional protein expression using two sequential approaches: 1) Per-sample detection of cancer cells using epithelial markers and clustering, 2) Annotation of stromal and immune populations using TRIBUS ^17^ - a prior knowledge-based tool that combines unsupervised clustering with highly granular hierarchical automated cluster annotation (**Fig. 1b to e**, **Supplementary Fig. 1b**). We comprehensively validated the cell phenotyping by visual assessment of the cell annotation masks overlayed with the highly multiplexed images via customized Napari plugin ^18^ (**Fig. 1d,e**). The deep cell phenotypes comprised two non-immune stromal populations (annotated stromal) constituting 40% of the cells (CD31+ endothelial cells and Vimentin+, α-SMA+ fibroblasts) and 15 immune populations corresponding to 17% of the cells, including B cells, memory and regulatory T cells, and Myeloid lineages such as macrophages, neutrophils and antigen-presenting cells (**Fig. 1f,g**). The per-sample abundance of CD8+ T cells and CD20+ B cells concord with the expert pathologists’ annotations based on IHC (**Supplementary Fig. 1c,d**). The annotated cell phenotypes also showed agreement when projected on UMAPs. For example, lymphocytes (B cells and T cells) formed distinct clusters in the UMAP projection, while Myeloid cells depicted a phenotypically more heterogeneous group (**Fig. 1g**). After cell phenotyping, we quantified the functional cell states (**Fig. 1h**) and cell-to-cell spatial neighborhoods (**Fig. 1i**). To decode the phenotypic and spatial heterogeneity of cancer cells ^19^, we clustered the cancer cells based on the expression of cancer-cell phenotypic markers, mesenchymal and functional markers, and cell morphology features. Using unsupervised consensus clustering (see Methods), we uncovered four distinct cancer cell phenotypic populations (**Supplementary Fig. 1e**) with distinct molecular and morphological characteristics (**Fig. 1k, l; Ext. Figure 1, 2**).

### Spatial TME is shaped by molecular profiles and chemotherapy exposure

We set out to evaluate clinically relevant patterns of immune infiltration across the defined clinicomolecular groups. We observed increased immune cell infiltration in BRCAloss and HRD tumors across both PDS and IDS samples (**Supplementary Fig. 1f**), with a notable enrichment of CD4+ T cells. Interestingly, lymphocyte populations demonstrated significant co-infiltration patterns within the TME (**Supplementary Fig. 1g**). Furthermore, the abundance of several lymphocyte populations was associated with longer progression-free survival (PFS) (**Supplementary Fig. 1h**) and overall survival (OS) (**Supplementary Fig. 1i**). Notably, in IDS samples, most immune cell infiltrations lacked prognostic significance, suggesting that tumor-intrinsic factors contribute to prognosis ^14,20^. Additionally, the highly intercorrelated infiltration patterns limit the interpretability of individual hazard ratios, and imply that lymphocyte infiltration represents a coordinated process potentially orchestrated through shared biological mechanisms.

To map the spatial TME domains we clustered the 4,644,485 single cells based on their spatial neighborhood characteristics (see Methods). We uncovered 14 phenotypically distinct spatial clusters, hereafter referred to as recurrent cellular neighborhoods (RCN) (**Fig. 1m,n, Supplementary Fig. 1j**). Four types of spatial domains were identified containing stromal (RCN01 to RNC03), cancer (RCN04 to RNC08), immune and stromal cells adjacent to cancer cells (annotated tumor-stroma interface, TSI; RCN09 and RCN10) and immune neighborhoods (RCN11 to RCN14). The RCN abundances were highly intercorrelated (**Supplementary Fig. 1k**) indicating that the TME domains are spatially co-orchestrated. The cancer spatial domains included a heterogenous phenotypic neighborhood (RCN04), and four neighbourhoods predominated by distinct tumor cell functional phenotypes. Interestingly, the majority of cancer cells were localized in the RCN4 neighborhood containing the most heterogeneous cancer cell phenotypic composition measured using Shannon diversity index, indicating the preferred topographical concurrence of heterogeneous tumor cell clones (**Fig. 1m**). Further, the RCN4 was less abundant in the IDS samples (**Fig. 1o,p**) suggesting phenotypic convergence of cancer cell populations following chemotherapy, consistent with our previous observation ^14^. Moreover, we observed significant variations in the abundance of the immune-cold stroma (RNC01), enriched in the CCNE1 and HRP tumors, and immune-cold TSI (RCN09) enriched in HRD-IDS tumors (**Fig. 1o**). Comparison of the spatial domains across therapy sequences further showed enrichment of stromal-non endothelial neighborhoods (RCN01 and RCN02), and immune-cold TSI (RCN09) in the chemotherapy exposed IDS samples, and proliferative tumor cell neighborhood (RCN7) in the PDS samples (**Fig. 1p**). These results highlight that the spatial TME domain topographies are shaped by the molecular profiles and chemotherapy exposure.

### Spatial tumor-stroma interface shapes clinical outcomes

Among all the RCNs, the immune-infiltrated TSI (RNC10) exhibited the highest diversity (Shannon diversity index: 2.3, **Fig. 1m**), exhibiting the most heterogeneous cell composition among all the RCNs. Statistical comparison of the two TSI types, revealed that immune-infiltrated TSI contained significantly higher proportions of T cells, myeloid cells, macrophages, and B cells suggesting the presence of co-orchestrated immune infiltration (**Fig. 1q**). Consistent with our previous observations ^6^, the BRCAloss tumors contained the highest abundance of immune-infiltrated TSI (**Supplementary Fig. 1l**). By contrast, the immune-cold TSI (RCN9), which was more abundant in the IDS samples compared to PDS (**Fig 1o**), was significantly enriched in fibroblast stromal cells (**Fig 1q**). Additionally, the immune-cold TSI showed a higher prevalence of ‘Idle’-type cancer cells lacking activity-linked phenotypic marker expressions, indicative of an ‘immune-evasive’ TSI where cancer cells could evade the recognition of the immune system (**Fig. 1q**). Conversely, the immune-infiltrated TSI (RCN10) had significantly more active cancer phenotypes, with enriched proliferative, immunogenic and EMT phenotypes suggestive of adaptive “immune escape” mechanisms at the TSI ^21^.

Next, we assessed which RCNs were most strongly associated with clinical outcomes. Using adjusted Cox regression models stratified by therapy sequence (PDS/IDS), we identified three RCNs significantly associated with both PFS and OS (**Supplementary Fig. 1m, n**). Notably, the TSI spatial domain and its multicellular composition emerged as a key determinant of OS in multivariable Cox models across the entire patient cohort (**Fig. 1r**). Specifically, a high abundance of the immune-infiltrated TSI (RCN10) was associated with improved prognosis, whereas the abundance of immune-cold TSI (RCN09) correlated with poorer outcomes (**Fig. 1r**).

These findings underscore the critical role of multicellular spatial domains and immune cell infiltrations in shaping the clinical outcomes within the HGSC TME. Among these, the TSI emerges as the most diverse spatial domain, exhibiting divergent phenotypic compositions indicative of unique mechanisms underlying immune tolerance. The intricate interplay between immune co-infiltration, spatial organization, and clinicomolecular characteristics highlights a coordinated orchestration of these processes. This emphasizes the necessity of integrative approaches to uncover the drivers of the observed spatial phenotypes within the HGSC TME.

### CEFIIRA enables the discovery of clinically relevant single-cell spatial tumor ecosystem features

To reveal the biological processes underlying spatial immune infiltration patterns and prognosis, we developed CEFIIRA (Cell Feature Importance Identification by RAndom forest), a robust and generalized machine learning (ML) framework (**Fig. 2a**). The primary objective of CEFIIRA is to identify the most distinguishing features within high-dimensional single-cell data that differentiate between two sample groups. CEFIIRA robustly minimizes bias in both ML model creation and the identification of critical features by: 1) Using ML model parameter tuning and feature importance evaluation in cross-validation, training and testing the model on multiple data partitions to prevent overfitting, 2) Ensuring that both the training and test sets have a proportional and balanced representation of samples from each class, 3) Validation of ML models on a prior defined validation set with unseen data, 4) Partitioning single-cell data in a sample-wise manner to minimize data leakage, 5) A robust identification of critical features using feature elimination and feature permutations, 6) Performing the previous steps over 30 iterations (nested cross validations) to achieve a consensus estimation. Additionally, our versatile, open-source software CEFIIRA can integrate multi-level data from each sample, such as clinical information from the patient and molecular profiles (**Fig. 2a**).

**Figure 2.**
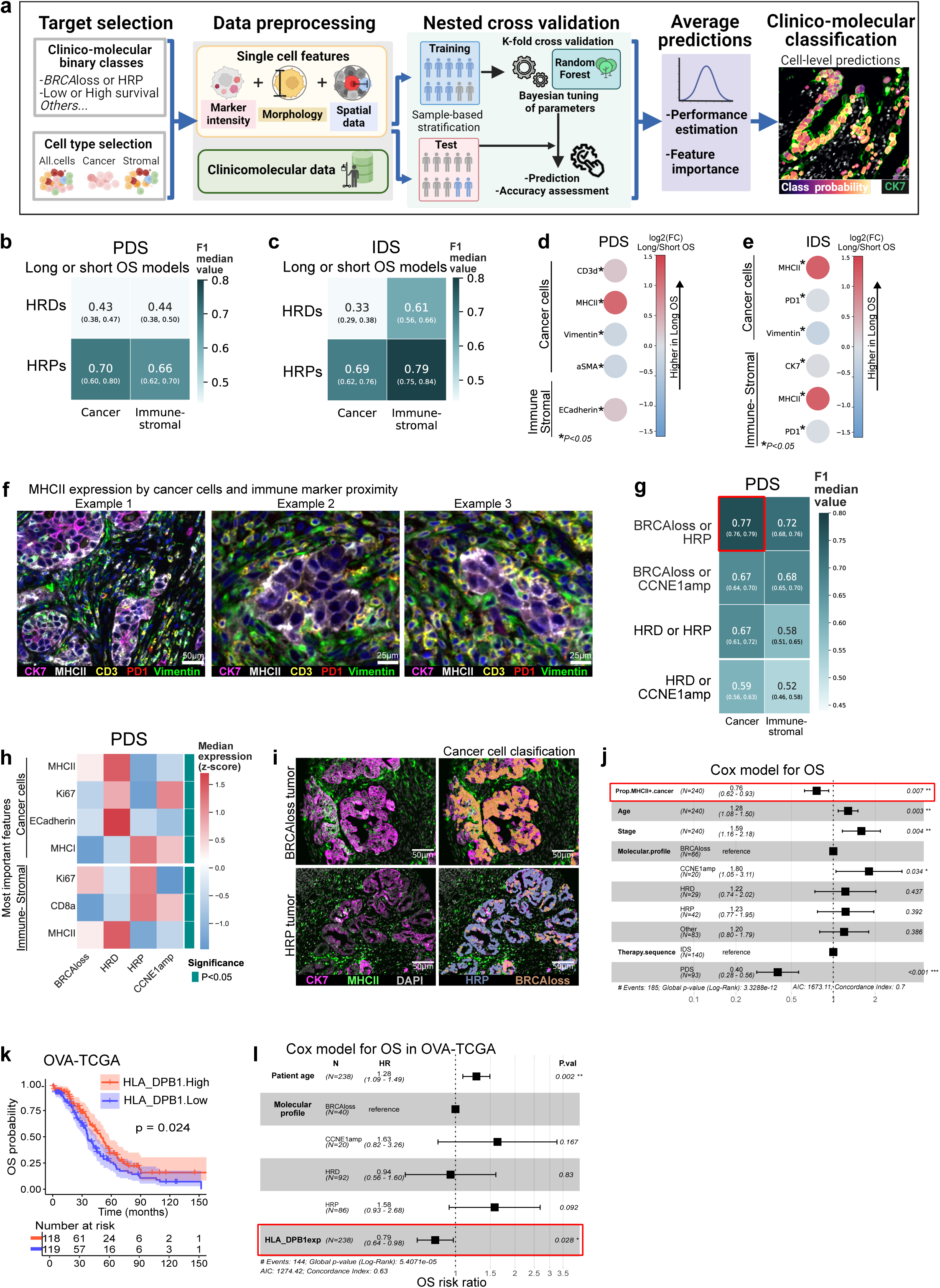
**a** A schematic illustrating our machine learning pipeline ‘CEFIIRA’. Firstly, we defined targets for binary predictions, for example, prediction if cancer cells belong to either BRCAloss or HRP tumors. Data preprocessing is performed for the tumors related to the target (in the example, BRCAloss and HRP). Next, 30 nested cross-validation iterations are performed. In each iteration, 80% of the tumors (their corresponding cells) are randomly assigned to the training set, with the remaining 20% allocated to the test set. Subsequently, a random forest model is built using the training set, with several parameters optimized through Bayesian tuning and K-fold cross-validation. The trained model is validated on the test set to predict the target classes (in the example, whether cancer cells belong to either BRCAloss or HRP tumors) and its most predictive features are calculated. After completing 30 iterations of nested cross-validations, the model’s median accuracy and consensus critical features are identified. **b-c** Heatmap displaying the median accuracy (F1 score) of CEFIIRA models in predicting whether cells belong to tumors from short- or long-survival patients, across the indicated molecular profiles (rows) and cellular populations (columns). The 95% confidence intervals for the F1 scores are shown in parentheses. The results corresponding for PDS samples are in panel **b** and in panel **c** for IDS samples. **d-e** Dot plots displaying the log2 fold change in marker intensity between cells from short- and long-survival patients. The markers displayed correspond to the top five most important features for each model (excluding clinical variables). Only markers that were statistically different between short- or long-survival groups are shown (Wilcoxon rank sum test, unadjusted p < 0.05). The assessment was performed for HRP tumors, the analysis for PDS samples are shown in panel **d** and in panel **e** for IDS samples. **f** tCycif images of independent samples, displaying the expression of relevant markers. **g** Heatmap of the median accuracy prediction (F1 score) of the indicated targets (rows) when using different cellular populations (columns). The corresponding models were constructed and validated using PDS samples. The 95% confidence intervals of the F1 scores are shown in parentheses. **h** Heatmap displaying the median expression, by molecular profile, of the top five most important features shared by models predicting: ‘BRCAloss or HRP’, ‘BRCAloss or CCNE1amp’, and ‘HRP or HRD’. The selection of important shared features was done independently for cancer cells and immune-stromal cells. The median expression shown was quantified in the specified cell populations. Their differential expression across the molecular profiles was assessed using a two-tailed Kruskal-Wallis test; the significant differences (unadjusted p<0.05) are highlighted in teal boxes. **i** tCycIF images of a BRCAloss tumor in the upper panels and of a HRP tumor in the bottom. Cancer-cell class predictions for BRCAloss and HRP are displayed in the right panels. **j** Forest plot of the Cox regression of overall survival (OS) using the indicated variables and including the proportion of MHCII+ cancer cells per tumor (prop.MHCII+.cancer). **k** Kaplan-Meier plot displays the overall survival (OS) probability associated with the deconvoluted cancer cells-specific RNA expression of the MHCII gene HLA_DPB1. The cohort corresponds to HGSCs from OVA-TCGA. The stratification of tumors as High and Low expression was according to the median value across them. Statistical significance was assessed using a Log-rank test. **l** Forest plot of the Cox regression of overall survival (OS) using the indicated variables and the deconvoluted cancer cells-specific RNA expression of the MHCII gene HLA_DPB1 in OVA-TCGA.

We aggregated the multidimensional spatial data for each of the 4,644,485 single cells, including the phenotype and functional marker expressions (**Fig. 1d, h**), morphological features and spatial neighborhoods (**Fig. 1i**), resulting in 43 feature representations per cell (Materials and Methods). To uncover multicellular features that differentiate the TME in terms of OS, we stratified the patients into three equally sized groups: long, medium, and short OS (**Supplementary Fig. 2 a, b**). We generated CEFIIRA models for either cancer cells or the immune-stromal compartment separately in HRD-like (BRCAloss, HRD) or HRP-like (HRP, CCNE1amp), and in PDS or IDS tumors respectively, also including age and tumor stage as potential confounders. As expected, the patient’s age and tumor stage were the top predictive features of long-OS for most of the models (**Supplementary Fig. 2c,d**). The models trained on HRP-like tumors exhibited better classification performance than those trained on HRD-like tumors in both PDS (**Fig. 2b**) and IDS (**Fig. 2c**), indicating that CEFIIRA can separate the patients with differential prognosis within the HRP-like tumors using multimodal single-cell spatial data.

### Cancer cell intrinsic MHCII expression is a key determinant of overall survival

Within the HRP tumors, CEFIIRA identified cancer-cell intrinsic MCHII as one of the key features for long or short OS prediction in both PDS and IDS samples (**Fig. 2d, e, Supplementary Fig. 2c,d**). MHCII was also the most enriched feature in tumors from long-OS patients compared to short-OS patients (**Fig. 2d, e**). By contrast, stromal markers Vimentin, α-SMA in PDS samples, and immune checkpoint receptor PD-1 in IDS samples were enriched in tumors with shorter OS. In addition, CD3d expression was a predictive marker of long-OS in the PDS samples (**Fig. 2d**). The prevalence of stromal and immune markers in cancer cell models suggested the relevance of signals from neighboring cells in the TSI. Indeed, visual inspection of the tCycIF images confirmed MHCII expression by cancer cells, and that the Vimentin, CD3d, and PD-1 signals were primarily signals originating from distinct neighboring cells adjacent to the cancer cells (**Fig. 2f**). In addition, MHCII was also the top predictive marker in the immune-stromal cells of IDS samples (**Fig. 2e)**. Altogether, CEFIIRA models revealed that MHCII and immune-stromal markers, indicative of cancer-stromal spatial proximity, are critical for OS prediction regardless of therapy sequence.

Next, to determine if MCHII expression is specific to a molecular profile, we trained CEFIIRA models to predict the molecular profile of individual cells. We observed that PDS samples demonstrated superior classification performance than IDS samples (**Fig. 2g**, **Supplementary Fig. 2f**). In cancer cell models, MHCII was a key feature with higher expression in the HRD-like tumors (**Fig. 2h, Supplementary Fig. 2g**). The CEFIIRA model to classify BRCAloss or HRP cells achieved the highest predictive performance (Median F1=0.77, **Fig. 2g**), with MHCII being the most differential feature displaying higher expression in BRCAloss cancer cells (**Supplementary Fig. 2g, h**), also confirmed from the tCycIF images (**Fig. 2i**). These results indicate that cancer intrinsic MHCII expression is enriched in the HRD tumors.

To validate the predictive role of cancer-intrinsic MHCII expression on prognosis across different molecular profiles, we performed an adjusted Cox regression on the entire cohort, including the tumors not assigned to a particular molecular profile and not used in the CEFIIRA analysis. In the entire cohort, a higher proportion of MHCII+ cancer cells consistently predicted longer OS (**Fig. 2j**) further strengthening the link between MHCII expression in cancer cells and prolonged OS in HGSC, regardless of the molecular profiles or therapy sequence. To further corroborate the prognostic role of MHCII expression by cancer cells in a larger dataset, we extended our validation to The Cancer Genome Atlas (TCGA), by inferring the cancer-cell specific MHCII expression from bulk RNA-seq using PRISM ^22^. Consistently, we observed that high MHCII expression by cancer cells predicted better OS in this independent cohort at mRNA level (**Fig. 2k, l**). These findings confirm that MHCII expression in cancer cells is a predictor marker of improved clinical outcomes in HGSC, suggesting a link to distinct tumor-immune interactions within the TME.

### Cancer MHCII expression shapes immune infiltration

To investigate the effect of cancer-intrinsic MHCII expression on overall immune infiltration, we compared the abundance of immune populations according to the fraction of MHCII+ cancer cells per tumor. This analysis revealed that a higher fraction of MHCII+ tumor cells associated with enhanced infiltration of CD4+ and CD8+ T cells, CD15+ myeloid cells, and M1-like macrophages (CD11c+ or CD207c+) (**Fig. 3a**). To corroborate these findings, we tested the association between MHCII expression in cancer cells and the estimated proportions of immune cells using deconvoluted data from the TCGA (CIBERSORT, see Methods). We found that tumors with high MHCII expression in cancer cells were significantly enriched with T cells and M1-like macrophages (**Supplementary Fig. 3a**). In contrast, tumors with high MHCII expression had the lowest signal of immunosuppressive M2-like macrophages, suggesting a more favorable immune microenvironment for anti-tumor responses (**Supplementary Fig. 3a**). Overall, these results demonstrate that HGSC tumors with a high abundance of MHCII+ cancer cells have a more immune infiltration in the TME.

**Figure 3.**
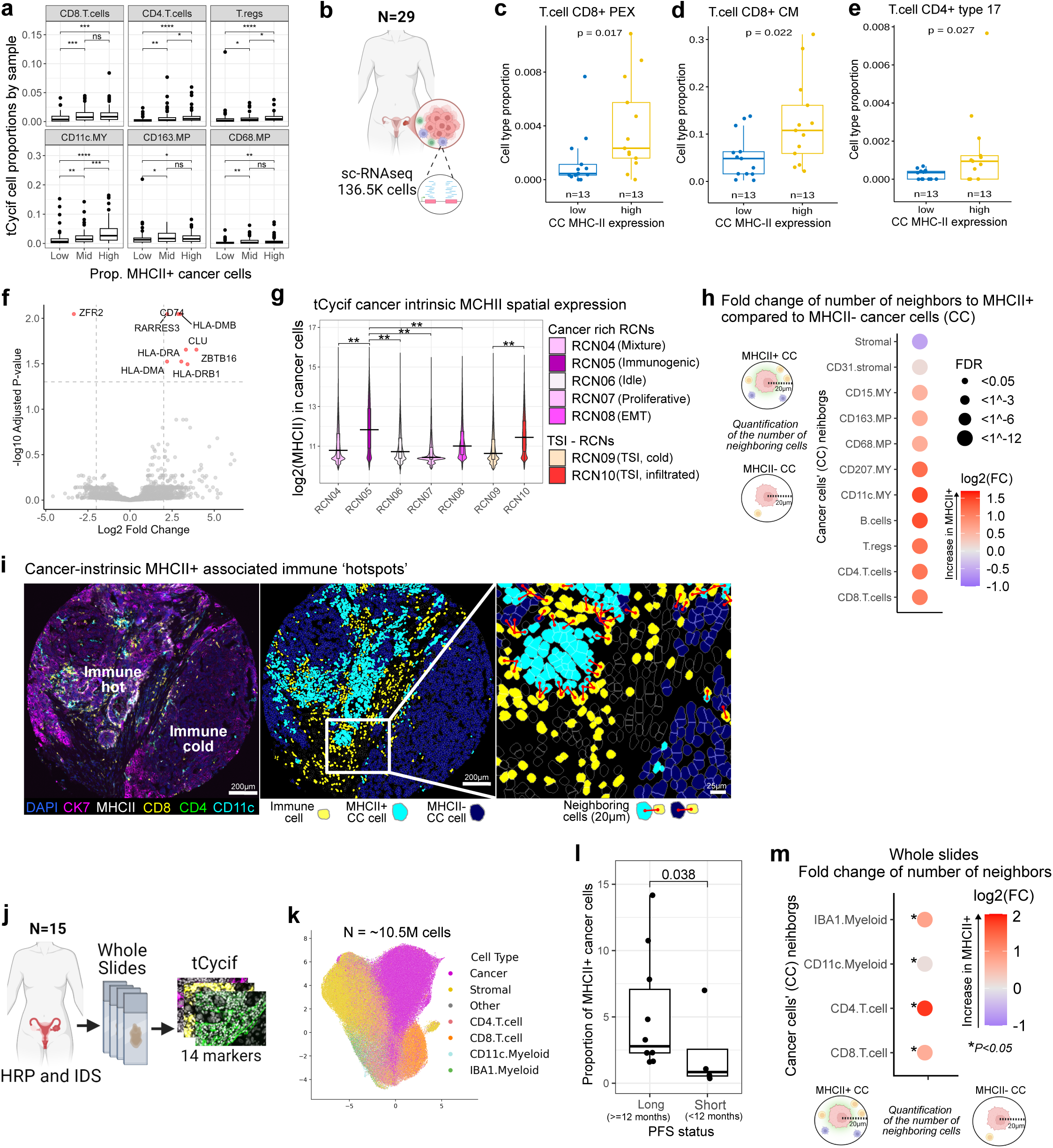
**a** Boxplots depicting the proportions of immune populations in the tCycif images from the multi-omics cohort. The tumors are grouped in three equal-sized groups, according to the fraction of MHCII+ cancer cells. In the boxplots, black horizontal lines represent the median, the boxes extend from the first to the third quartile and the whiskers indicate the values at 1.5 times the interquartile range. Outliers are shown as dots. The differences in proportion were calculated using a two-tailed Wilcoxon rank-sum test. The notation for P-values are as follow: *P<=0.05, **P<=0.01, ***P<=0.001, ****P<=0.0001. **b** Schematic of the sc-RNAseq dataset analyzed. **c to e**, Boxplots depicting the estimated proportions of immune populations per tumor analyzed by sc-RNAseq, with low and high number of MHCII+ cancer cells. Panel **c** for CD4+ type 17 helper T cells, **d** for CD8+ precursor exhausted cells (PEX) T cells, and **e** for CD8+ central memory (CM) T cells. The differences in proportion were calculated using a two-tailed Wilcoxon rank-sum test. In the boxplots, black horizontal lines represent the median, the boxes extend from the first to the third quartile and the whiskers indicate the values at 1.5 times the interquartile range. **f** Volcano plot of the pseudo-bulk differential expression analysis in cancer cells, comparing samples with a high proportion of MHCII+ cancer cells with those with low MHCII+ cancer cells. **g** Violin plots of the expression of MHCII (logarithmic base 2 transformation) in cancer cells within the indicated Recurrent Cellular Neighborhoods (RCNs). In pink are displayed the RNCs enriched in cancer cells, and in orange and yellow RNCs localized in the Tumor-Stroma Interface (TSI). The differences between profiles were calculated using a two-tailed Wilcoxon rank-sum test. The black horizontal lines represent the median, the boxes extend from the first to the third quartile and the whiskers indicate the values at 1.5 times the interquartile range. **h** Dotplot displaying the log2 Fold Change (log2FC) in the number of neighbors to MCHII+ cancer cells compared to MHCII- cancer cells on the entire multi-omics cohort. Only dots representing statistically significant differences are shown (Wilcoxon rank sum test, FDR < 0.05). A panel is shown for PDS and IDS samples. **i** tCycIF image and graphical representation displaying intra-tumor MHCII expression in cancer cells and immune cell proximity. The immune ‘hot’ and immune ‘cold’ areas were manually annotated according to the abundance of immune cells observed in them. **j** Schematic of the whole slide image analysis in a collection of 15 HGSC samples. All the tumors were HRP and were extracted from IDS. **k** UMAP projection of cells from the whole slide images, the colors correspond to their main cell type group assignment. **l** Boxplot depicting the proportion of MHCII+ cancer cells per tumor. The patients were stratified by progression-free survival (PFS) time, where PFS-long corresponds to more than 12 months of PFS, otherwise was considered PFS-short. The abundance difference was calculated using a two-tailed Wilcoxon rank-sum test. The black horizontal lines represent the median, the boxes extend from the first to the third quartile and the whiskers indicate the values at 1.5 times the interquartile range. Individual points represent tumors. **m** Dotplot displaying the log2 Fold Change (log2FC) in the number of neighbors to MCHII+ cancer cells compared to MHCII- cancer cells. Only dots representing statistically significant differences are shown (Wilcoxon rank sum test, unadjusted P < 0.05).

To further investigate the distinct immune cell proportions and underlying transcriptional programs, we analyzed 51 HGSC samples, both from IDS and PDS, from 29 patients using single-cell RNA sequencing^14^ (sc-RNAseq) (**Fig. 3b**). The dataset comprised 136,452 cells, which included 25,091 epithelial/cancer cells, 13,952 non-immune stromal cells, and 97,409 immune cells, annotated using canonical phenotypic markers as described previously ^14^ (see Methods). Using gene signature analysis, 12% of cancer cells had a high MHCII expression level similar to that of canonical antigen-presenting cells (**Supplementary Fig. 3b**). We observed that tumors with a high proportion of MHCII+ cancer cells had increased proportions of type 17 proinflammatory helper CD4+ T cells ^23^, central memory (CM) CD8+ T cells, related to activation after antigen exposure ^24,25^, and precursor exhausted (PEX) CD8+ T cells, which have traits of both exhaustion and memory ^26^ (**Fig. 3c to e**). Interestingly, pseudo-bulk differential expression analysis on cancer cells, revealed that tumors with a high proportion of MHCII+ cancer cells had upregulated genes related to cellular stress response and apoptosis (**Fig. 3f**). Altogether, these findings suggest that MHCII+ cancer cells influence immune infiltration by promoting antigen presentation and driving proinflammatory responses.

### Cancer cell MHCII expression is spatially coordinated and associated with immune cell proximity

To assess the relationship between MHCII expression and the spatial organization of the TME, we analyzed cancer-intrinsic MHCII expression across the cancer cell spatial domains (RCN04 to RNC08). Our analysis revealed high levels of MHCII expression within RCN05 **(Fig. 3g**), abundant in cancer cells co-expressing other immunogenic markers, including MHCI, pSTAT1, pTBK1, PDL1, (**Fig. 1k**). This indicates that MHCII+ cancer cells are often spatially aggregated even within the cancer cell spatial compartments.

We next explored the relationship between the cancer cell MHCII expression and the cell composition at the TSI. Comparison of the two TSI spatial domains, revealed a significantly higher expression of MHCII in cancer cells within the immune-infiltrated TSI (RCN10) as compared to the immune-cold TSI (RCN9) (**Fig. 3g**). Consistently, analysis of the cell-cell physical proximity revealed that MHCII+ cancer cells resided more often in close proximity to immune cells as compared to MCHII- cancer cells across the diverse clinicomolecular profiles (**Fig. 3h, Supplementary Fig. 3c**). In contrast, MHCII- cancer cells were predominantly neighboring stromal cells. Visual inspection of the tumor tissues confirmed that MHCII+ cells were surrounded by dense clusters of immune cells, forming immune ‘hotspots’, while MHCII- cells were embedded in immune ‘cold’ tumor niches (**Fig. 3i**).

To expand our analyses to whole tissue sections, we generated high-dimensional single-cell spatial data using tCycIF on 15 HRP-IDS samples (**Fig. 3j**) from an independent cohort in the Oncosys-OVA prospective trial (NCT06117384). We captured and phenotyped 10.5 million cells using the TRIBUS software (Fig. 3 k**, Supplementary Fig. 3d**). Consistently, a higher proportion of MHCII+ cancer cells associated with longer progression-free survival (PFS) (**Fig. 3l**). By contrast, CD8+ T-cell infiltration did not associate with PFS (**Supplementary Fig. 3e**), in agreement with the observation in the multi-omics cohort, underscoring the presence of tumor-intrinsic factors influencing prognosis in IDS samples. Importantly, we observed that the MHCII+ cancer cells (**Supplementary Fig. 3d**) had significantly more neighboring CD4+ T, CD8+ T, and CD11C+ Myeloid cells as compared to the MHCII- cells, generating spatial immune hotspots (**Fig. 3m**). Altogether, these findings demonstrate that MHCII+ cancer cells colocalize in immunoreactive spatial domains, driving immune infiltration, tumor-immune interactions and shaping the TSI.

### Cancer-intrinsic MHCII governs single-cell and spatial immune activation

Next, we set out to uncover transcriptional programs that underpin the immune infiltration induced by the MHCII+ cancer cells. Interestingly, tumors with a high MCHII expression exhibited upregulated genes related to immune activation including chemokines, cytokines, immune checkpoints, granzymes, and immune receptors, compared to tumors with low MCHII expression in the TCGA dataset ^27^ (**Supplementary Fig. 4a, b, Supplementary Table 3**). Notably, we observed a significantly augmented expression of the immune checkpoint molecules TIGIT, CTLA4 and TMIGD2 in the MCHII+ tumors. Gene overrepresentation analysis ^28^ revealed 14 enriched pathways related to immune activation (**Fig. 4a**), including both PD-1 and chemokine signaling, and IFN-γ activation. No pathways were enriched within the under-expressed genes. Consistently, a high proportion of MHCII+ cancer cells was associated with overexpression of chemokines, cytokines, immune checkpoints, and immune receptors in the Nanostring data of the multi-omics cohort (**Supplementary Fig. 4c, Supplementary Table 4**). Interestingly, CXCL9, CXCL10, and CXCL11 chemokines, and their receptor CXCR3, associated with interferon-gamma (IFN-γ) activation ^29–31^, were among the top overexpressed genes in both cohorts, suggesting that IFN activation plays a key role in MHCII driven immune activation as previously suggested also in other cancers ^32–34^.

**Figure 4.**
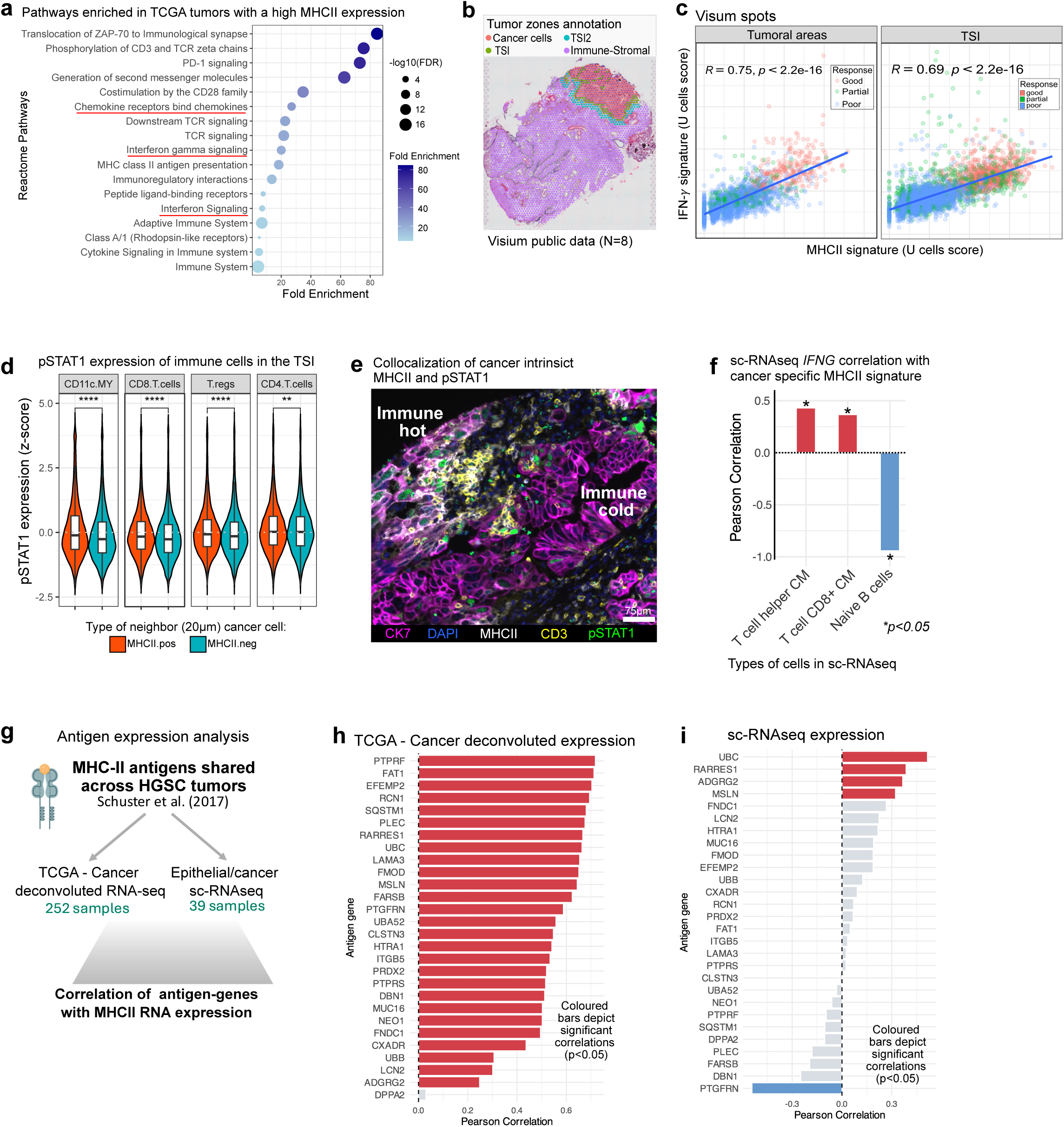
**a** Pathways overrepresented (enriched) based on the overexpressed genes when comparing samples with low vs high cancer-deconvoluted MHCII in TCGA. The overrepresentation analysis was performed using the PANTHER web tool, and the statistical significance was assessed using Fisher’s exact test. **b** Tumor zones annotations of expert pathologists from matching H&E slides of Visium spatial transcriptomic data. **c** Scatter plot displays the MHCII and interferon gamma (IFN-γ) U-cell signatures in the Visum spots from the zones annotated as ‘tumor’ and ‘TSI’. Statistical significance was assessed using Pearson correlation. The spots are colored according to the patient’s response to chemotherapy. **d** Violin plots depict the pSTAT1 expression in immune cells neighboring cancer cells in a radius of 30μm. The violins are grouped according to the MHCII signal status (positive or negative) of the neighboring cancer cell. The expression difference was calculated using a two-tailed Wilcoxon rank-sum test. The black horizontal lines represent the median, the boxes extend from the first to the third quartile and the whiskers indicate the values at 1.5 times the interquartile range. The notation for P-values are as follow: *P<=0.05, **P<=0.01, ***P<=0.001, ****P<=0.0001. **e** tCycIF image displaying an expression of MHCII and pSTAT and cell type markers. The immune ‘hot’ and immune ‘cold’ areas were manually annotated according to the abundance of immune cells observed in them. **f** Barplots displaying the Person’s correlation of *INFG* gene expression in the listed immune cell types, with MCHII signature expression in epithelial/cancer cells, inter tumors using pseudo-bulk estimations. Only significant associations are shown (p<0.05). **g** Schematics of the analysis of the association between antigen expression and MHCII expression in cancer cells. **h and i** Barplot displaying Pearson’s correlation between the expression of HGSC antigen-genes reported by *Schuster, H et al 2017*, and MHCII expression by cancer cells (deconvoluted component) in TCGA cohort (**b**) and sc-RNAseq dataset using pseudo-bulk estimations (**c**).

To explore the spatial correlations between MHCII expression and IFN-γ signaling, we conducted an in-depth re-analysis of a previously available spatial transcriptomic dataset ^35^ incorporating expert pathologist annotations and spatially resolved signature analyses (**Fig. 4b**, **Supplementary Fig. 4d**). The annotated TME areas exhibited distinct transcriptomic profiles depicting differential clusters in a UMAP projection (**Supplementary Fig. 4e**). Using MHCII and IFN-γ signatures ^36^ (see Methods), we found that the MHCII expression was significantly correlated with IFN-γ signature expression across distinct tumor areas, including cancer and TSI regions (**Fig. 4c**). Moreover, a higher MHCII signature and increased FN-γ in both cancer and TSI spatial regions predicted improved clinical responses to chemotherapy (**Supplementary Fig. 4f, g**). Interestingly, in our tCycIF spatial data (multi-omics cohort), statistical analysis revealed significantly higher expression of phospho-STAT1 (pSTAT1) indicative IFN activation in CD8+ and CD4+ T cells and antigen-presenting (CD11c+) myeloid cells neighboring MHCII+ cancer cells at the TSI (**Fig. 4d**), confirmed via visual inspection (**Fig. 4e, Supplementary Fig. 4h**). These findings suggest that cancer cell MHCII associates with spatially coordinated activation of antigen presentation and T cells via IFN signaling at the TSI. Further, Using the sc-RNAseq dataset, we observed that the MHCII expression in cancer cells correlated with *INFG* gene expression specifically in CD4+ memory helper and CD8+ central memory (CM) T cells, which are related to activation after antigen exposure (**Fig. 4f**). Overall, these findings reinforce the link between cancer intrinsic MHCII and IFN-γ activation and highlight the role of cancer cell MHCII as a driver of spatial immune activation.

### Cancer intrinsic MHCII is associated with antigen expression

Canonically, MHC-II complexes are stable on the cell surface only when antigen-loaded, and typically present exogenous antigens or endogenous misfolded proteins^37^. To investigate whether cancer-specific MHCII expression is linked with specific antigen presentation, we focused on previously detected antigens using immuno-peptidomics in HGSC ^38^ (**Fig. 4g**). We found that the MHCII expression in cancer cells indeed correlated with the expression of most antigen-encoding genes in the TCGA dataset (**Fig. 4h**). However, in our sc-RNAseq dataset, only four specific antigen genes (*UBC*, *RARRES1*, *ADRG2*, *MSLN*) were expressed in the cancer cells, and correlated significantly with MHCII expression (**Fig. 4i**). These results suggest that cancer cell-intrinsic antigen presentation via MHCII is an active process and underlying spatial immunogenicity contributing to T-cell activation in HGSC. Enhancing MHCII antigen presentation directly by cancer cells could boost immunogenicity and open new avenues for immunotherapeutic interventions in HGSC.

## DISCUSSION

We constructed a spatial atlas of the HGSC tumor ecosystem using 929 comprehensive maps of the multicellular TME derived from 265 real-world clinical samples. Our spatial analysis paradigm identified distinct spatial domains with varying phenotypic compositions. These intercorrelated domains were linked to specific tumor cell phenotypic profiles and clinicomolecular groups, underscoring the role of molecular drivers and chemotherapy in shaping TME topographies. Notably, of the two distinct TSI phenotypes, the immune-infiltrated TSI was associated with cancer cell phenotypic heterogeneity and improved outcomes, while the immune-cold TSI displayed stromal predominance and was linked to poorer outcomes. Using our machine learning framework CEFIIRA we integrate high-dimensional single-cell data and uncovered MHCII as a key prognostic marker in HGSC. Through spatial profiling of ∼15.1 million cells across two independent cohorts, we show that MHCII+ cancer cells within the immunogenic tumor and TSI neighborhoods generate immune hotspots. The hotspots showed evidence of coordinated immune activity with enhanced cellular interactions of MHCII+ cancer cells particularly with T cells and antigen-presenting myeloid cells. Interestingly, CD4+ T cells have been found to recognize MHCII+ cancer cell antigens in melanoma and breast cancer-promoting immune surveillance ^39,40^, and the spatial contact between MHCII+ cancer cells and CD8+ T cells was reported recently in triple-negative breast cancer ^41^. While IFN expression in tumors has traditionally been linked to immune surveillance, recent evidence suggests it may also play a role in immune escape mechanisms ^42^. Through a robust spatial TME analysis, we found that MHCII+ cancer cells preferentially cluster spatially within immune-reactive cancer-cell neighborhoods and the TSI with augmented IFN activity. Moreover, the cancer cell MCII correlated with IFN produced by antigen experienced CD4+ and CD8+ memory T cells, implying that the MCII tumor cells can actively present antigens and activate the local TME. Furthermore, we identified that MHCII and *IFN-*γ expression are spatially correlated, and demonstrated cancer intrinsic MHCII expression induces IFN activation of neighboring CD4+ and CD8+ T, and antigen-presenting myeloid cells at the TSI. These findings, summarized in **Fig. 5**, support the hypothesis that via distinct antigen presentation, the tumor-intrinsic MHCII activates CD4+ and CD8+ T cells to produce IFN resulting in augmented immune activation at the TSI associating with favorable prognosis in HGSC.

**Figure 5.**
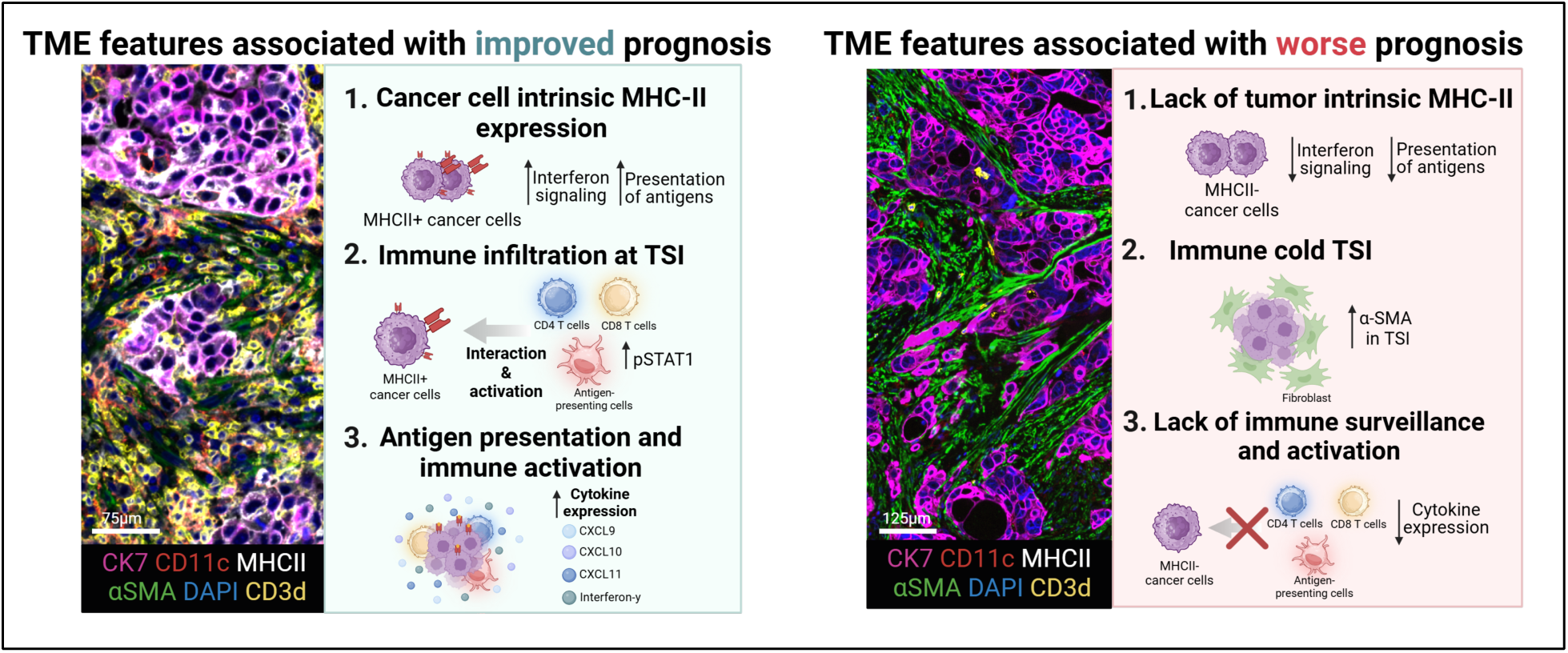
Illustration highlighting the main findings in our study.

The presence of MHCII+ tumor cells has been linked to an improved response to immune checkpoint inhibitors in melanoma, lung cancer, and triple-negative breast cancer ^33,41,43,44^. This suggests that MHCII expression might similarly influence the efficacy of immunotherapies in HGSC, however, only modest benefits have been observed in clinical trials using single-agent PD-1/PD-L1 inhibitors in unselected patient populations ^45^. Our study revealed that other immune checkpoint inhibitors, such as TIGIT and CTLA4, were overexpressed in tumors with high MHCII expression, emphasizing the need for further clinical exploration of immune checkpoint inhibitors beyond PD-1/PDL-1 in selected patients with HGSC. While tumor-intrinsic MHCII expression indicates a functional tumor-immunity cycle via active antigen presentation, other therapeutic approaches should be developed for patients with tumors lacking MHCII expression. In conclusion, our single-cell spatial atlas uncovers mechanisms orchestrating immune surveillance across the clinicomolecular subtypes of HGSC providing rationale for future therapeutic approaches and patient stratification.

### Study limitations

We acknowledge certain limitations in our study highlighting the opportunities for future refinement and exploration. The retrospective design and reliance on archival tissues may introduce biases related to sample preservation and heterogeneity. This was managed through careful quality control, although a subset of tumors was excluded from downstream analysis to ensure data reliability. Our protein marker panel, while comprehensive, has inherent limitations in representing all relevant cell types. While our transcriptomic findings were validated in independent single-cell datasets, there is potential to enhance the accuracy of deconvolution methods. For our machine learning models, incorporating robust approaches to address correlated traits could further improve predictive performance. Lastly, our analysis culminated in a testable hypothesis, which we encourage future studies to explore through functional models to deepen understanding.

## ACKNOWLEDGMENTS

We acknowledge the NKI-AVL Core Facility Molecular Pathology & Biobanking for supplying NKI-AVL Biobank material and lab support. This work was supported by the Dutch Cancer Society [IKNL2014-6838]. This study was co-funded by the European Union (ERC, SPACE 101076096). Views and opinions expressed are however those of the authors only and do not necessarily reflect those of the European Union or the European Research Council. Neither the European Union nor the granting authority can be held responsible for them. This study was also funded by the Sigrid Jusélius Foundation, Research Council of Finland (grant numbers 1339805, 350396), Cancer Foundation Finland, University of Helsinki Research Foundation (F.P.), Ida Montinin Säätiö (F.P.), The Päivikki and Sakari Sohlberg Foundation (F.P), Orion Research Foundation (F.P.), Biomedicum Helsinki Foundation (F.P.). We also wish to thank the IT Center for Science (CSC) for the computational resources, Angela Szabo for figure design suggestions, and Ali Amiryousefi for suggestions in the development of machine learning models.

## ETHICS APPROVAL

For the present study, approval of the institutional review boards of the Netherlands Cancer Registry (NCR), PALGA, and NKI-AVL was obtained for the 265 patient cohorts. Informed patient consent was waived according to Dutch legislation due to the use of archival samples and pseudonymization of the material. The whole slide tumor specimens were collected as part of the ONCOSYS-OVA study (NCT06117384) in accordance with the ethical standards from the 1975 Declaration of Helsinki. Each patient gave their written informed consent and the use of samples and clinical data was approved by the local Ethics Committee (HUS334/2021).

## AUTHOR CONTRIBUTIONS

F.P. performed study conceptualization and design, gathering of resources, data curation, image analysis, statistical analysis, interpretation of results, guided the machine learning pipeline development, and drafting of the manuscript. L.v.W. performed gathering of resources, data curation, clinical data analysis guidance and drafting of the manuscript. A.S. performed machine learning analysis, data analysis, interpretation of the results, and drafting of the manuscript. A.J. performed gathering of resources and data analysis. Z. K. performed image data analysis. I.N. performed spatial transcriptomics data analysis. M.M.F. performed sc-RNAseq data analysis.

E.A. performed image data curation. J.C. performed image analysis and provided guidance. E.M. guided the machine learning pipeline development. E.K. performed pathologist annotations of Visium data. F.C. performed image data curation. M.S. performed clinical data curation. S.S. performed gathering of resources. S.R., J.C., M.G., performed data analysis and guidance for antigen analysis, G.S.S., M.A.vd.A., K.K.Vd.V, and R.FPM.K contributed to reviewing, editing and gathering of resources. A.Vi. performed pathologist annotations. V.C., A.Vä., and P.K.S. provided resources and guidance. A.F. and H.M.H. performed study conceptualization and design, interpretation of results, drafting of the manuscript, and supervision.

## DATA AVAILABILITY

Fluorescence microscopy images and cell feature quantification will be provided upon reasonable request to the corresponding author for non-commercial and academic purposes, and all requests will be promptly reviewed within 15 working days. Anonymous clinical data sharing from the NCR will be considered for non-commercial, research, or statistical-based use on a case-by-case basis (to be requested and approved by the NCR). The human sequence data generated in this study are not publicly available due to patient privacy requirements but are available upon reasonable request from the corresponding author.

## MATERIALS & METHODS

### Cohort description

We collected tumor samples from 265 HGSC patients with FIGO stage IIb-IV tumors from the Netherlands Cancer Institute - Antoni van Leeuwenhoek Hospital (NKI-AVL), Maastricht University Medical Centre (MUMC) and Amsterdam University Medical Centre (AUMC)), between January 2008 and December 2015. The patients were treated with primary cytoreductive surgery (PDS) or with neoadjuvant chemotherapy followed by interval debulking surgery (IDS). All patients received adjuvant platinum-based chemotherapy and paclitaxel, and none of them received Bevacizumab or PARP inhibitors as maintenance therapy. Clinical data was extracted from the Netherlands Cancer Registry (NCR). Histopathological data and Formalin-fixed, paraffin-embedded (FFPE) tissue blocks were obtained from the nationwide network of the Dutch Pathology Registry (PALGA) ^46^.

Tissue blocks originated from tumors retrieved during debulking (PDS or IDS) surgery. The histology of all tumors was confirmed by three dedicated pathologists (KvdV, HMH). Of the samples, 70% were extracted from primary tumors (ovaries, tuba or adnexal sites). FFPE blocks were organized into single tissue microarray (TMA) with one-millimeter diameter cores and with an average of four cores per patient’s sample. Immunohistochemical (IHC) staining was done on all cores to assess CD8+, CD20+, CD68+, and CD103+ cell expression as previously described^47^.

### Whole slide cohort description

We utilized HGSC tumors from 15 patients participating in the ONCOSYS-Ova clinical trial (NCT06117384). The samples were obtained from patients during interval debulking surgeries (IDS) conducted at Helsinki University Hospital, Finland (HUS334/2021). The selection criteria for these samples included HR-proficient status, a tumor purity greater than 10%, patients exhibiting a progression-free survival (PFS) of more than 6 months and a tumor macroscopic residual disease (R > 0cm) after surgery. The median PFS for patients with HRP tumors that underwent IDS therapy sequence is 11.7 months in the entire ONCOSYS-Ova clinical trial; based on that we selected a cutoff of 12 months to define long-PFS (>=12 months) and short-PFS (<12 months) in the survival analysis.

### Molecular profile assignment

For the TMA cohort, we identified *BRCA1/2* mutation and *BRCA1* promotor methylation status of each sample as previously described ^47^. Next, tumors without a *BRCA1/2* mutation or *BRCA1* promotor hypermethylation underwent shallow whole genome sequencing (sWGS) using Illumina HiSeq2500 to determine a *BRCA*-like CNV profile ^48^ and CCNE1-amplification status as described previously ^47^. In addition, using the sWGS, we performed HRD status estimation using the software shallowHRD ^49^. Finally, we categorized the tumors as *BRCAloss*, HRD, HRP and CCNE1amp using a hierarchical approach described in **Supplementary Figure 1a**.

For the TCGA, we annotated the tumors as HRD and HRP based on ovaHRDscar ^50^. For all analyses, only tumors with a *TP53* mutation, histological grades G3 and G4, and those at high stages (stage III and stage IV) were included. For the whole slide cohort, we adopted the clinically validated HRD classification performed at the Helsinki University Hospital, Finland which uses ovaHRDscar ^50^, and the detection of somatic and germline mutations and gene deletions in *BRCA1/2* genes.

### Highly multiplexed immunofluorescence and image preprocessing

To facilitate throughput and minimize experimental batch effects, tumor tissues were represented in tissue microarrays (TMA). To capture the whole tumor ecosystem, we sampled two cores from the tumor center and the tumor-stroma interface, resulting in a total of 1026 tissue spots. The TMAs were stained with a validated 30 plex antibody panel (**Supplementary Table 2**) and scanned with a RareCyte CyteFinder scanner following the tCycIF protocol ^51^. Illumination was corrected using the BaSiC tool ^52^. The per-image tiles were stitched and registered using ASHLAR (https://github.com/labsyspharm/ashlar). Individual TMA’s cores were cropped using the Image Processing Toolbox on MATLAB (version 2018b, The MathWorks, Inc., Natick, Massachusetts, United States). Cell segmentation was performed using Ilastik v1.3.2 ^53^ and Cellprofiler v3.1.8 ^54^ using 2-pixel nuclear expansion and the median intensity of each channel and morphological features such as Area, Eccentricity, Solidity, Perimeter, and Roundness were quantified using MATLAB. Quality filtering was applied based on changes in the DNA channel intensity across cycles (https://github.com/yunguan-wang/cycif_analysis_suite), image regions with artifacts, and TMA’s cores (n=64) with less than 1000 high-quality cells.

For the whole slide validation dataset, FFPE tissue sections were collected from 15 tumors. The slides were stained with a 14-plex validated antibody panel (**Supplementary Table 5**) and scanned with a RareCyte CyteFinder scanner following the tCycIF protocol. The image tiles were stitched with ASHLAR. The single-cell objects were segmented using nuclei segmentation in StarDist ^55^ implementing the pre-trained model ‘*2D_versatile_flu*o’’. The mean intensity of each channel was quantified by our in-house script (https://github.com/farkkilab/image_processing/tree/main) based on skimage package ^56^. Image quality control was performed by CyLinter ^57^.

### Cell phenotyping

All cell phenotypes were curated via visual inspection. For the TMAs cell phenotyping, we first classified cells as either cancer or non-cancer on a per-sample basis. Cells were clustered based on the Z-score scaled signal of epithelial markers (Cytokeratin 7 and E-cadherin) using a Gaussian mixture model from the R-package Mclust. Two Gaussian clusters were fit, and cells in the higher epithelial marker expression cluster were labeled as cancer cells. Clustering was manually reviewed per sample, and poorly performing samples (n=10) were excluded. Non-cancer cell annotation was performed for each TMA slide. Outliers above the 99.9th percentile and below the 0.01th percentile were adjusted to these values. Each channel’s signal was Z-score scaled, and non-cancer cells were clustered by marker intensity (**Supplementary Fig. 1c**) into a 2500-node self-organized map (SOM) using the R-package flowSOM. For each SOM node, we computed cell-type label scores using the flowSOM function QueryStarPlot, first identifying global cell types (non-immune stromal and immune cells), and then annotating immune subtypes (**Supplementary Fig. 1b**). After annotation, we quantified cell population proportions per TMA core and estimated the median proportion for each patient’s sample.

For whole slide images, the cell type annotation was performed using our software Tribus ^17^. Outliers were removed using 99.9th percentile for input marker intensity, normalized via arcsin transformation with a cofactor of 5 ^58^, and scaled by Z-score. Tribus was run with the following parameter settings of *sigma=1*, *learning_rate=1*, *clustering_threshold=100*, *undefined_threshold=0.0005*, *other_threshold=0.4*. A total of 6 cell types were annotated including tumor, stromal, CD4+ T-cell, CD8+ T-cell, CD11c+ myeloid and IBA1+myeloid. We used a constant logic table (**Supplementary Table 6**) for all images when running Tribus. The exceptions are images S100_iOme and S110_iOme, which were manually gated with Scimap due to nonspecific staining CD8a in cancer cells. We corroborated our annotations by comparing them to cancer cell proportions determined by expert pathologist annotations using hematoxylin and eosin (H&E) staining, showing agreement across samples (**Supplementary Fig. 3g**). The MHCII+ cancer cells were identified using manual thresholds and comparing the annotations to the tCycIF images.

### Cancer cell phenotype clustering

Firstly, we scaled the marker signals by TMA slide and performed outlier-cell signal truncation, as mentioned in the previous method section. We perform dimensionality reduction the cancer cells according to the marker expressions: “Ki67”, “Vimentin”, “pSTAT1”, “MHC-I”, “ECadherin”, “yH2AX”, “pTBK1”, “cPARP1”, “CK7”, “PDL1_488”, and morphological features: “Area”, “Eccentricity”, and “Roundness” into a 2500-node self-organized map (SOM) using the R-package flowSOM ^59^. The SOM-nodes were clustered using the R-package ConsensusClusterPlus ^60^. For this, the number of clusters (*k*) was assessed from 2 to 12. For each value of *k* assessed, 500 iterations were performed, in each of the 80% of the SOM-nodes and 90% of the features (markers and morphological features) were randomly selected. The algorithm “partitioning around medoids” (pam) was used to cluster the selected dataset per iteration. After the iterations, we determined the best number of cancer meta-clusters, *k*, by assessing the resultant consensus-clustering matrix for each value of *k* and its cumulative density distribution (**Extended Figure 1**). The cancer meta-clusters were named according to the expression of relevant markers in each one (**Extended Figure 2**).

### Recurrent cellular neighborhoods and spatial analysis

For each cell from the collection of 265 tumors, we computed the fraction of neighboring cell types in a radius of 30µm, using the function tl.spatial_count of the package Scimap ^61^. We clustered the cells based on the neighborhood information to generate recurrent cellular neighborhoods (RCNs), by using k-means and a k**=**40 employing the tl.spatial_cluster function of the package Scimap. To select the value of k, we used the function KElbowVisualizer of the Python package YellowBrick (**Extended Figure 3a**). We used a radius of 30µm to identify recurrent neighborhoods of cells that are in physical contact. The replication of our analysis using a radius of 60µm and 90µm led to similar clusters, which shows the robustness of our approach (**Extended Figure 3b**).

### Spatial transcriptomics analysis

To perform the analysis of the MHCII signature activity in the proximity of the tumor, we utilized a publicly available Visium spatial transcriptomics dataset, composed of 8 IDS HGSC samples with different responses to chemotherapy (GSE211956). For spatial gene expression data, we performed the QC and preprocessing of input feature-barcode matrices in h5 format, using the Seurat R package ^62^. For the QC, we excluded from the analysis the spots with less than 500 genes detected, more than 25% of mitochondrial genes or more than 20% of hemoglobin genes. We normalized the feature-barcode matrices using Seurat’s SCTransform function.

Based on H&E histological images, each spot was manually annotated by an expert pathologist as ‘Cancer’, ‘TSI’ (tumor-stroma interface with partial tumor and stromal morphology), ‘Immune-Stromal’ and ‘other’. Spots annotated as ‘other’ have been excluded from the analysis. Within ‘Immune-Stromal’ spots, we have distinguished an additional type - ‘TSI2’, defined as Immune-Stromal spots with the closest proximity to either ‘Cancer’ or ‘TSI’ spots. Next, we assessed the activity of MHCII- signature across the spot types, by computing its normalized UCell score, using the R UCell package ^36^. The MHCII signature comprised of the following genes: *HLA-DMA*, *HLA-DMB*, *HLA-DOA*, *HLA-DOB*, *HLA-DPA1*, *HLA-DPB1*, *HLA-DQA1*, *HLA-DQA2*, *HLA-DQB1*, *HLA-DRA*, *HLA-DRB1*, *HLA-DRB5*, *CD74*. Finally, we’ve compared the distribution of the computed MHCII-signature UCell scores across different spot types, regarding the response to chemotherapy. The categorization of tumors as good, partial or poor response was adopted from Böhm et al, 2015 ^63^. Statistical significance has been computed using a two-sided Wilcoxon Rank Sum test.

### Machine learning supervised classification

The input to models consisted of single-cell data, where each cell had the following pre-processed features: (1) log2-transformed protein marker expression data, scaled by slide using standard scaler (z-score); (2) three morphological features (area, eccentricity, and roundness), scaled using standard scaler; (3) 14 features for the proportion of neighbors in a 30μm radius, one feature per annotated cell type; (4) patients’ clinical information like FIGO stage and age when the target was clinical outcome like short- or long-OS. For each analysis, cells with marker and morphological feature values outside the 0.01 to 0.99 quantile range were excluded. In addition, samples were excluded if they had less than 100 Cancer, or Immune-Stromal cells. For the creation of the subsequent machine learning models, to prevent group leakage, stratified sampling was employed, ensuring patients were divided 80% into training and 20% into the test sets maintaining balance across classes.

To identify the most suitable supervised machine learning (ML) algorithm for classifying single cells into molecular profiles, we used the PyCaret library on the entire dataset (https://github.com/pycaret/pycaret). We employed different ML algorithms for multiclass predictions of: *BRCA*loss, HRD, HRP, and CCNE1amp. Based on PyCaret runs, we selected Random Forest (RF) as a suitable algorithm based on its mean performance (weighted-averaged AUC and F1) on ten-fold cross-validations (**Supplementary Table 7**). In addition, we utilized the PyCaret RF model to determine the number of classes that lead to better prediction (weighted-average AUC and F1) using the same pre-processed single-cell dataset, running independent models for four, three and two classes using ten-fold cross-validations. Based on the results we opted to further construct models for two classes (**Supplementary Table 8**).

Based on our comparisons, we created an RF pipeline using the Python scikit-learn library ^64^ for binary classification of cells into clinicomolecular groups (https://github.com/farkkilab/CEFIIRA). The models’ training process employs nested cross-validation, where the outer loop iteratively partitions tumors into training and test sets 30 times using the stratified sampling previously mentioned. For each nested cross-validation, using the training set, a Bayesian optimization was used to tune the hyperparameters of the model (*min_samples_split*, *min_samples_leaf*, *n_estimators*). The Bayesian optimization process iteratively evaluates hyperparameters values’ combinations to find the ones that achieved the highest mean weighted F1 score across nine cross-validation iterations; for each iteration, the training set was divided into three-folds three independent times. Next, the created model is subjected to a cross-validated recursive feature elimination (RFECV) based on classification accuracy. This step enables identifying and extracting the most predictive features from the training set. We further assess the importance of these selected features in predicting the target variable through the computation of permutation scores. Finally, we evaluate the performance of the created model on the test set partitioned in the outer loop (unseen data) and extract single-cell level predictions of target classes. The median performance and standard deviation of the model’s in each of the 30 nested cross-validation is reported. Independent models were created for two cell compartments (Cancer cells, Immune-stromal cells). For the survival prediction models, we stratified the patients into long-survival groups (above the 66 percentile of overall survival time in the cohort) and short-survival (below the 33 percentile), independently for those with HRPs and HRDs tumors.

### Deconvoluted RNA-expression and CIBERT sort profile

For validation of the importance of immunogenic signals (MHCII and MHCI), we used publicly available RNA-expression data from The Cancer Genome Atlas (TCGA). The deconvoluted cancer-specific RNA expression was inferred with PRISM and adopted from previous work ^22^. The inferred cell proportions by CIBERSORT were obtained from the GDC-PanCanAtlas repository ^65^.

### RNA differential expression analysis and pathway enrichment

For the analysis of TCGA bulk RNA-seq, we obtained per sample gene read counts estimated with the software STAR ^66^, from the Genomics Data Commons (GDC). The associated clinical information was also obtained from GDC-PanCanAtlas repository ^65^. We stratified the tumors into three equal-size groups based on the expression of *HLA_DPB1* in the deconvoluted Epithelial Ovarian Cancer (EOC) component. We assessed the RNA differential expression between the tumors in the group with the highest (above quantile 0.66) and lowest (below quantile 0.33) expression of *HLA_DPB1* in the EOC component. We excluded the genes with less than 20 counts across the tumors from each assessed group. We used the DESeq2 R package ^27^ for count normalization and differential expression analysis. Genes were considered with differential expression according to the next criteria: Log2-fold change >=2, FDR < 1^-3. To identify over-represented pathways or sub-represented pathways we used as input the list of over-expressed or sub-expressed genes independently. Then, using the web portal PantherDB v19.0 ^28^, we performed a Fisher-exact test taking as reference the Reactome database.

For the analysis of RNA expression from Nanostring data. We first stratified the tumors into three equal-size groups according to the proportion of MHCII+ cancer cells using the tCycIF dataset. We assessed the differential expression between the tumors with the lowest (below quantile 0.33) and highest (above quantile 0.66) proportion of MHCII+ cancer cells. For the differential expression analysis, we computed per gene a two-sided T-test between the assessed groups, and the fold changes were estimated by calculating the difference in mean gene expression between the groups with the highest and lowest proportion of MHCII+ cancer cells.

Genes were considered with differential expression according to the next criteria: Log2-fold change >=1, FDR < 1^-3.

### sc-RNAseq analysis

We retrieved sc-RNAseq gene count matrices from 51 HGSC samples, pre-processed and annotated ^14^, from the Gene Expression Omnibus (GEO) database (accession code: GSE266577). Samples containing fewer than 40 epithelial cells were ignored from the analysis. We assessed the activity of MHCII signature across cell types by computing the normalized Ucell score using the R UCell package (v1.3.1) ^36^. The MHCII signature comprised of the following genes: *HLA-DMA*, *HLA-DMB*, *HLA-DOA*, *HLA-DOB*, *HLA-DPA1*, *HLA-DPB1*, *HLA-DQA1*, *HLA-DQA2*, *HLA-DQB1*, *HLA-DRA*, *HLA-DRB1*, *HLA-DRB5*, *CD74.* Based on the MHCII Ucell scores, the cells were stratified into three equal-sized groups (no expression, low expression, positive MHCII expression). According to the number of MHCII positive epithelial cells, the tumors were stratified into three equal-sized groups: high, medium, and low, according to tertile distribution. For each sample and celltype, pseudo-bulk gene expression profiles were generated using the function ‘*AggregateExpression’* from the Seurat package (v4.3.0)^67^. To assess the correlation of cancer-specific MHCII with IFN-γ in immune populations, it was computed a pseudo-bulk MHCII signature per sample using the previously mentioned list of genes and compared to the pseudo-bulk expression of the gene *IFNG* for each cell-type per sample. Only immune cells were selected for correlation analysis of MHCII- signature vs *IFNG* because they had a higher expression of *IFNG* as compared to Fibroblast or Epithelial cells (**Supplementary Fig. 4i**). Correlation was assesed using Pearson’s correlation.

### Survival analysis

To create survival plots and perform Log-rank tests and Cox regression models, we used the R packages survminer and survival. For TMAs, cell population abundance was estimated by calculating the median abundance per cell type across sampled cores. The proportion of MHCII+ cancer cells was determined relative to the total cancer cells per tumor. Tumors with at least 5% cancer cells and more than one core post-preprocessing were selected for survival analysis (n=234). Patient stratification based on cell population used the cohort’s median, defining high (above median) and low (below median) groups. For Cox models, tumors were split into three groups by abundance (below quantile 0.33, between 0.33-0.66, and above 0.66). Stratification by gene expression followed the same approach. Cox models were adjusted for age, FIGO stage, and molecular profiles where applicable. For TCGA dataset survival analysis, only high-stage HGSCs (III and IV) were included.

### Statistical analyses

We performed statistical analysis using R version 4.2.3. The difference in abundance between the two groups was calculated using a two-sided Wilcoxon rank sum test, unless stated that it was one-sided. The difference in abundance across different groups was calculated using a two-tailed Kruskal-Wallis test. The correlations between variables were assessed using Pearson correlations. For multiple testing, we adjusted P-values using the Benjamini and Hochberg method. *P*-values or adjusted P-values (FDR) less than 0.05 were considered statistically significant.

---

**Supplementary Figure 1**

**a** Assignment of the tumors into the four different molecular profiles. **b** Flowchart representing the strategy used to assign the cells from tCycIF images into different populations according to the expression of the indicated markers. **c,d** Boxplots of the proportion of cells in the tCycIF images, grouped by annotations made by an expert pathologist using immunohistochemistry (IHC). For each tumor, the pathologist’s annotation reflects the number of immune cells in the core with the highest number of IHC-positive cells (out of four cores per tumor), for example: <20 indicates fewer than 20 cells in the core with the most IHC-positive cells. **c** The panel displays the proportions of CD8+ T cells and panel **d** the proportion of CD20+ lymphocytes. The differences between groups were calculated using a two-tailed Wilcoxon rank-sum test. The black horizontal lines represent the median, the boxes extend from the first to the third quartile and the whiskers indicate the values at 1.5 times the interquartile range. Individual dots represent the values per tumor. **e** tCycIF image of the indicated markers, the right panel displays a magnification of the corresponding red area. Lower panel displays labels of the cancer cells according to their phenotypic cluster assignment. **f** Heatmaps of the median proportion of the indicated immune cells per molecular profile. A two-tailed Kruskal-Wallis test was used to calculate each immune cell type’s differential proportion across the molecular profiles. The significant differences (P<0.05) are highlighted in teal boxes. **g** Heatmap displaying the correlation of immune cell abundance across the tumors. Only pairwise significant (p<0.05) correlations are shown. **h** Forest plot of the Cox regression of progression-free survival (PFS) in relation to the abundance of the indicated immune cells per tumor. The regressions were adjusted by patient age, FIGO stage and molecular profile and were performed separately for PDS and IDS samples. The small squares represent the mean hazard ratio (HR), and the whiskers depict the 95% confidence intervals. **i** Forest plot of the Cox regression of overall survival (OS) in relation to the abundance of the indicated immune cells per tumor. The regressions were adjusted by patient age, FIGO stage and molecular profile and were performed separately for PDS and IDS samples. The forest plot description is as in panel h. **j** tCycif images with cell labels for the displayed RCNs. **k** Heatmap displaying the Pearson’s correlation for the abundance of each pair of RCNs across the samples. **l** Violin plots showing the proportion of RNC10 in PDS and IDS samples across molecular profiles. The differences between profiles were calculated using a two-tailed Wilcoxon rank-sum test. The black horizontal lines represent the median, the boxes extend from the first to the third quartile and the whiskers indicate the values at 1.5 times the interquartile range. Individual points represent tumors. **m, n** Forest plot of the Cox regression of overall survival (OS) for **m** and **n** for PFS, in relation to the abundance of the indicated RCN per tumor. The regressions were adjusted by patient age, FIGO stage and molecular profile and were performed separately for PDS and IDS samples. The small squares represent the mean hazard ratio (HR), and the whiskers depict the 95% confidence intervals. Only significant associations are shown.

**Supplementary Figure 2**

**a-b** Kaplan-Meier plots display the probability of survival of HRDs (BRCAloss and HRD) and HRPs (CCNE1amp, HRP) groups. The HRP and HRD groups are stratified using the 33 and 66 quantiles of survival times into long, medium and short overall survival (OS) times. The panel **a** displays PDS samples and panel **b** IDS samples. **c-d** A Heatmap displays the top seven important features for the models constructed to predict long or short-survival groups in HRDs or HRPs (columns). The features are grouped according to the cell populations used to create the models. The color values represent the Mean decrease in accuracy (MDA) when permuting the feature. Panel **c** corresponds to results when using PDS samples and panel **d** for IDS. **e** Violin plots depict the proportion of neighboring cells for the immune-stromal cells in the HRP tumors. The patients are grouped by short or long OS. The difference between groups was calculated using a two-tailed Wilcoxon rank-sum test. The notation for P-values is: ****P<=0.0001. **f** Heatmap of the median accuracy prediction (F1 score) of the indicated targets (rows) when using different cellular populations (columns). The corresponding models were constructed and validated using PDS samples. The 95% confidence intervals of the F1 scores are shown in parentheses. **g** Heatmap displaying the top five important features for the models constructed to predict the indicated targets (columns). The values represent the Mean decrease in accuracy (MDA) when permuting the feature. The targets are grouped according to the cell populations used for the models. **h** Radial plot depicts the ratio of median signal intensity of the indicated markers, between BRCAloss and HRP cancer cells.

**Supplementary Figure 3**

**a** Boxplots depicting the estimated proportions of immune populations in OVA-TCGA. The estimations were based on CIBERT sort analysis of RNA-seq data. The tumors are grouped in three equal-sized groups, according to the deconvoluted cancer cells-specific RNA expression of the MHCII gene HLA_DPB1. In the boxplots, black horizontal lines represent the median, the boxes extend from the first to the third quartile and the whiskers indicate the values at 1.5 times the interquartile range. Outliers are shown as dots. The differences in proportion were calculated using a two-tailed Wilcoxon rank-sum test. The notation for P-values are as follow: *P<=0.05, **P<=0.01, ***P<=0.001, ****P<=0.0001. **b** Violin plots showing the abundance of MHCII signature per cell population in the sc-RNAseq dataset. **c** Dotplot displaying the log2 Fold Change (log2FC) in the number of neighbors to MCHII+ cancer cells compared to MHCII- cancer cells across molecular profiles. Only dots representing statistically significant differences are shown (Wilcoxon rank sum test, FDR < 0.05). A panel is shown for PDS and IDS samples. **d** Flowchart representing the strategy used to assign the cells from whole slide tCycIF images into different populations according to the expression of the indicated markers. **e** Boxplot depicting the proportion of CD8+ T cells per whole slide images. The patients were stratified by progression-free survival (PFS) time, where PFS-long corresponds to more than 12 months of PFS, otherwise was considered PFS-short. The abundance difference was calculated using a two-tailed Wilcoxon rank-sum test. The black horizontal lines represent the median, the boxes extend from the first to the third quartile and the whiskers indicate the values at 1.5 times the interquartile range. Individual points represent tumors. **f** High-plex tCycIF images of a whole tissue slide, with magnifications to show areas containing MHCII+ cancer cells. **g** Scatter plot displaying the proportion of cancer cells identified in tCycIF images compared to the proportion annotated by expert pathologist using H&E images from adjacent slides.

**Supplementary Figure 4**

**a** Volcano plot displays differential gene expression results comparing tumors with high versus low cancer cells-specific RNA expression of the MHCII gene HLA_DPB1 in OVA-TCGA. The statistical inference was assessed using the DESEq2 package. In red are highlighted genes differentially expressed (log2-fold change >2, FDR < 1^- 3). **b** List of relevant genes differentially overexpressed from the analysis shown in panel b. **c** A volcano plot displays differential gene expression results comparing tumors with high versus low proportions of MHCII+ cancer cells in the multimodal cohort of 265 patients. The analyzed genes are from a custom gene panel, in which the *INFG* gene was not included. Statistical inference was calculated using a two-tailed Student’s t-test. In red are highlighted genes differentially expressed (log2-fold change >1, FDR < 1^-3). **d** Barplot depicting the number of Visium spots per annotated tumor zone in each of the eight analyzed tumors. **e** UMAP projection of Visum spots from all tumors based on their RNA expression. The colors represent the tumor zones to which they belong. **f,g** Violin plots show the MHCII signature for **f**, and **g** for IFN-γ signature abundance in the annotated tumor zones of the Visum dataset. The tumors are grouped according to the patient’s response to chemotherapy. Each dot on the violin plot represents a Visum spot. Its differential abundance across the associated response to chemotherapy was calculated using a two-tailed Wilcoxon rank-sum test. **h** tCycIF image displaying an expression of MHCII and pSTAT and cell type markers. The immune ‘hot’ and immune ‘cold’ areas were manually annotated according to the abundance of immune cells observed in them. **i** Pseud-bulk expression of *IFNG* across the annotated cell populations in the sc-RNAseq dataset. Each dot represents a tumor. The black horizontal lines represent the median, the boxes extend from the first to the third quartile and the whiskers indicate the values at 1.5 times the interquartile range. Outliers are shown in dots.

