## Supplementary figures and images for "Single-cell spatial atlas of high-grade serous ovarian cancer unveils MHC class II as a key driver of spatial tumor ecosystems and clinical outcomes"

### Extended Figure 2

**Extended material - Figure 2**

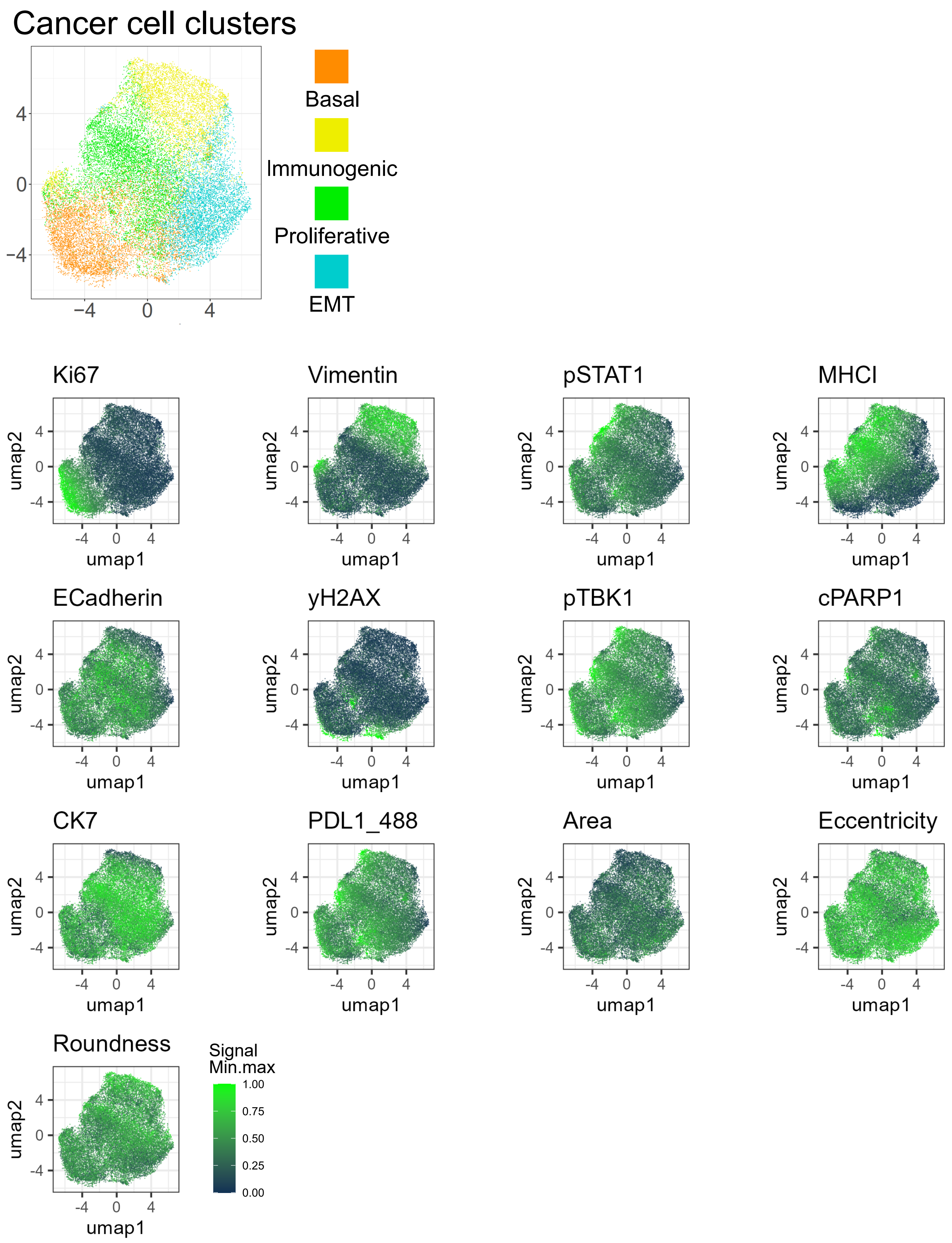

### Supplementary Figure 1

#### Supplementary Figure 1

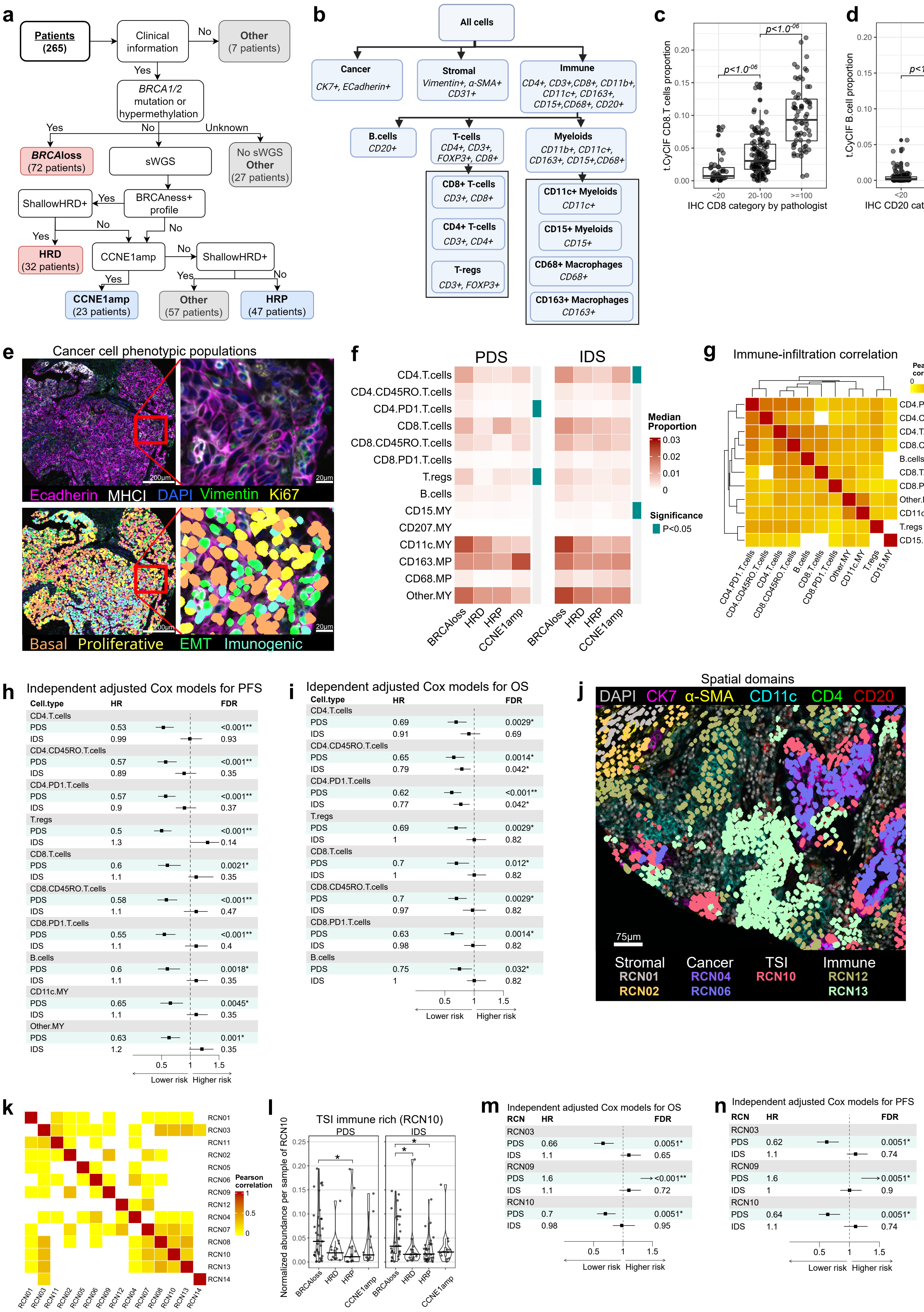

### Supplementary Figure 2

**Supplementary Figure 2**

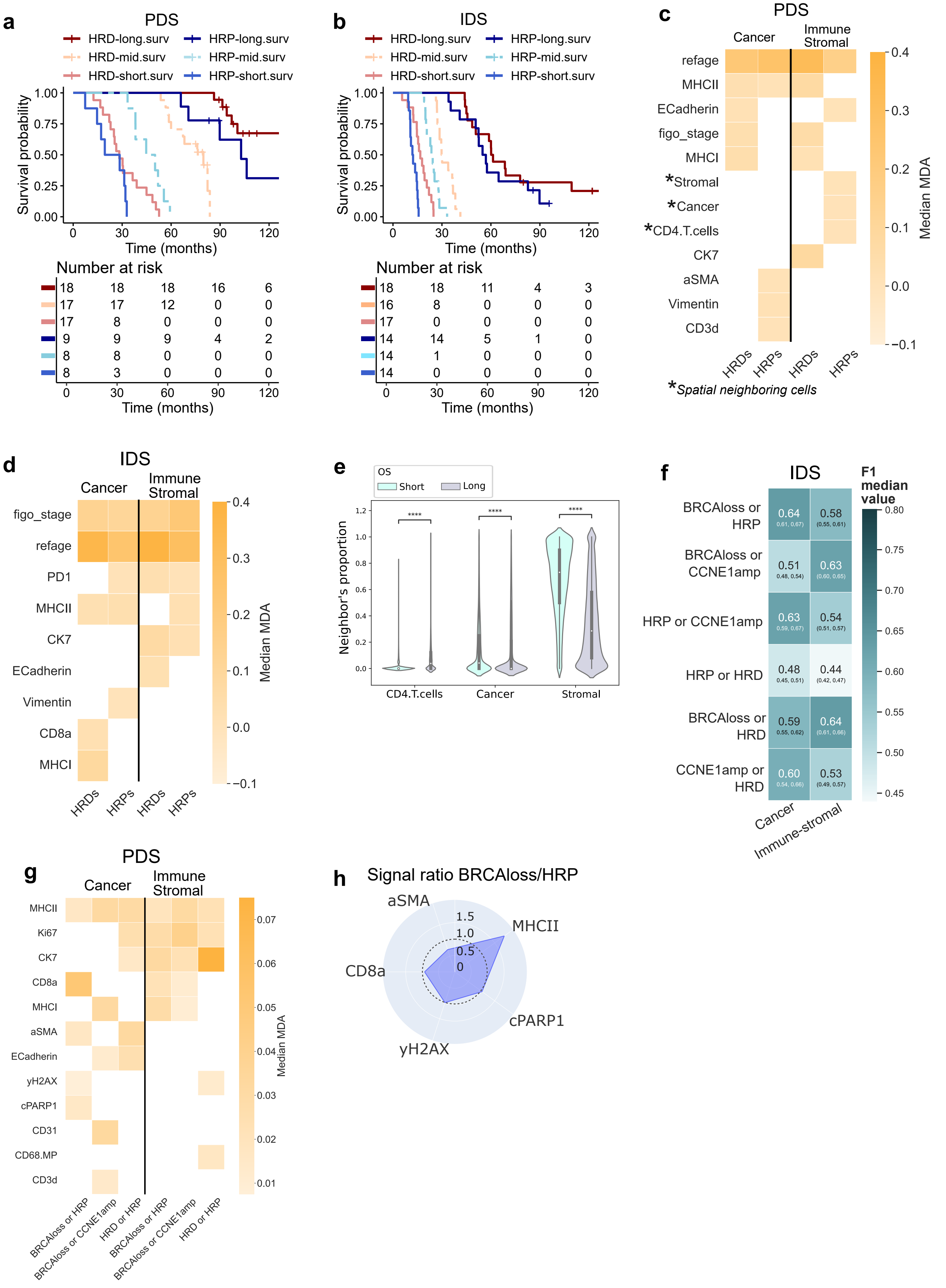

### Supplementary Figure 3

#### Supplementary Figure 3

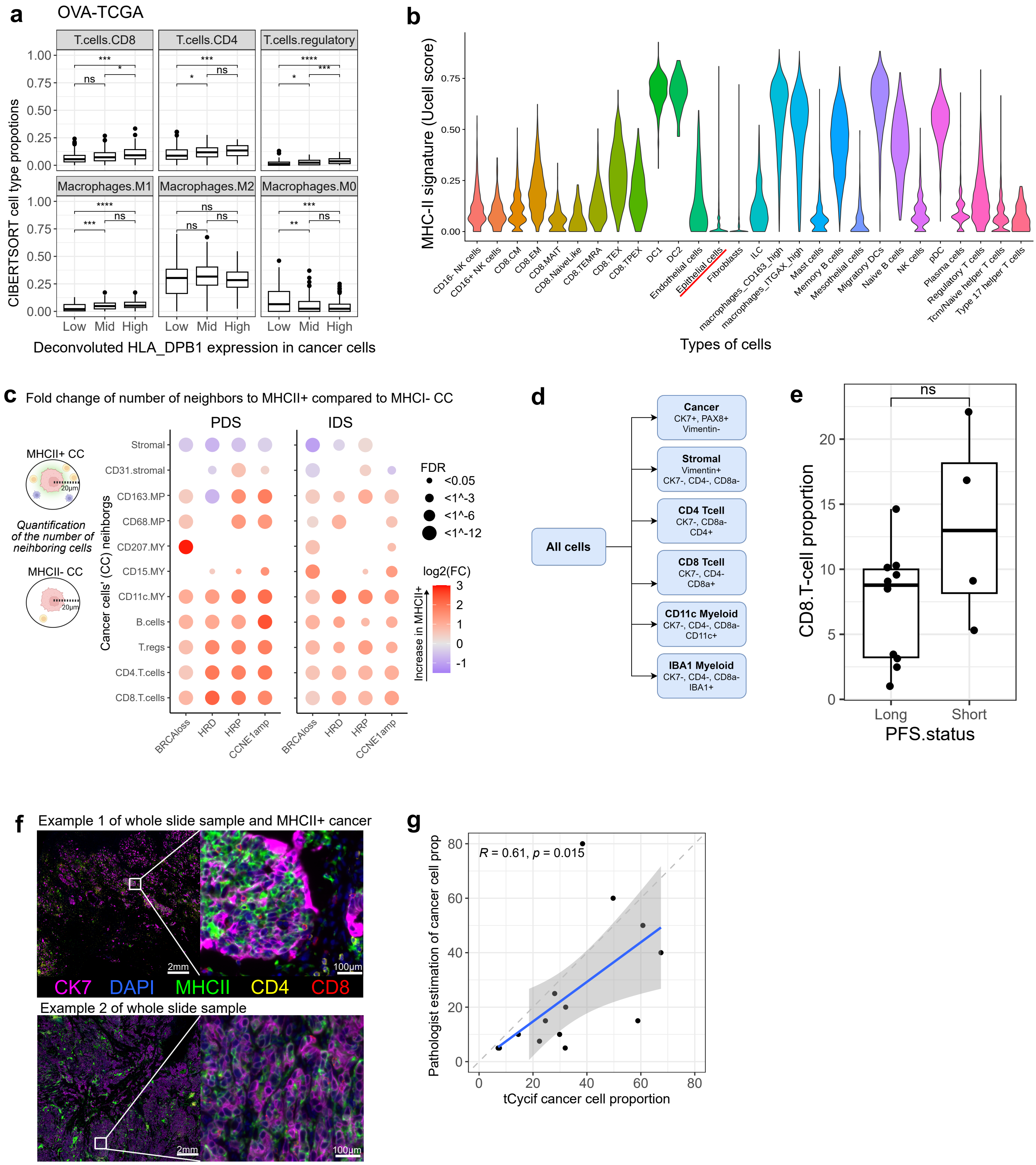
