## Supplementary Figure 4 for "Single-cell spatial atlas of high-grade serous ovarian cancer unveils MHC class II as a key driver of spatial tumor ecosystems and clinical outcomes"

a

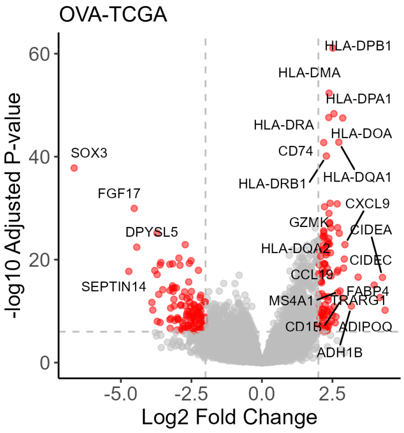


b

| Chemokines | Immune receptors |
|---|---|
| CCL5 | CD1A |
| CCL19 | CD1B |
| CCL22 | CD1C |
| CXCR3 | CD1E |
| CXCR6 | CD2 |
| CXCL9 | CD3D |
| CXCL13 | CD3E |
| XCL2 | CD8A |
| **Cytokines** | CD40LG |
| IL9 | CD244 |
| IL21 | CD247 |
| IL21R | FCRL1 |
| IFNG | FCRL4 |
| **Granzymes** | FCER1A |
| GNLY | FDCSP |
| GZMA | KLRC1 |
| GZMK | KLRC4 |
| GZMM | KLRB1 |
| **Immune checkpoints** | KIR2DL1 |
| CTLA4 | KIR2DL3 |
| TIGIT | PIGR |
| TMIGD2 | TRAT1 |

c

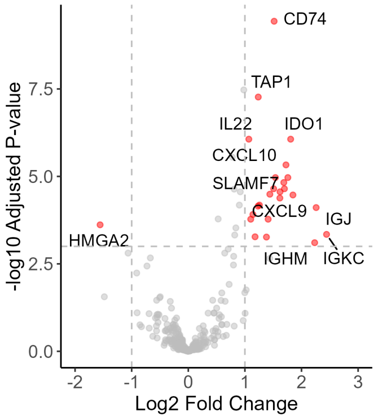


d

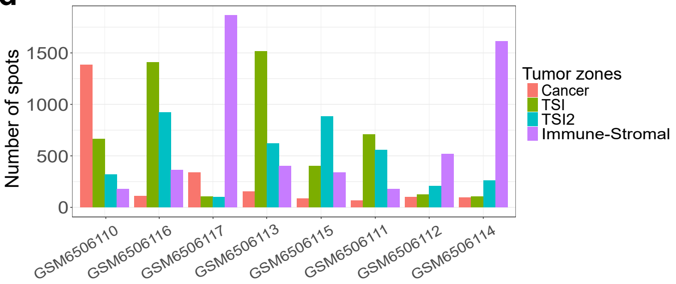


e

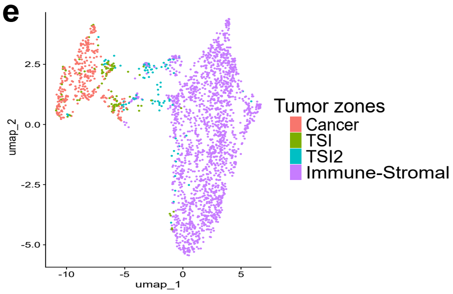


f

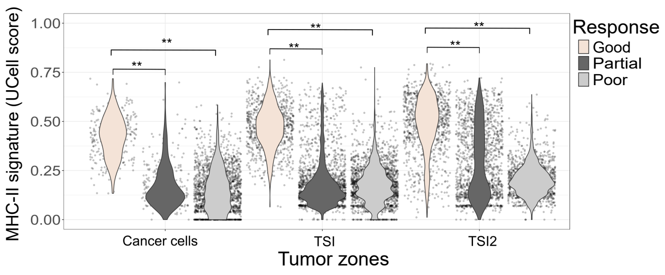


g

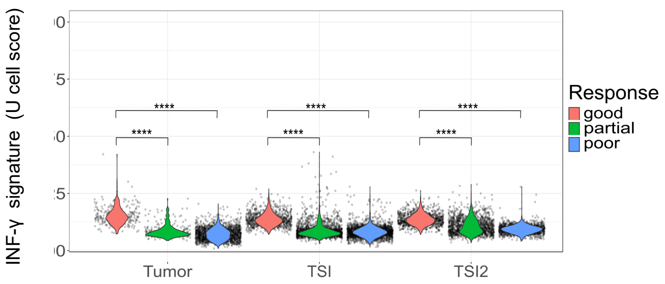


h

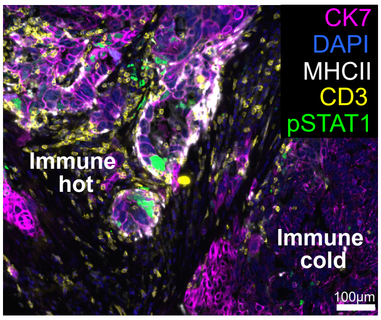


i

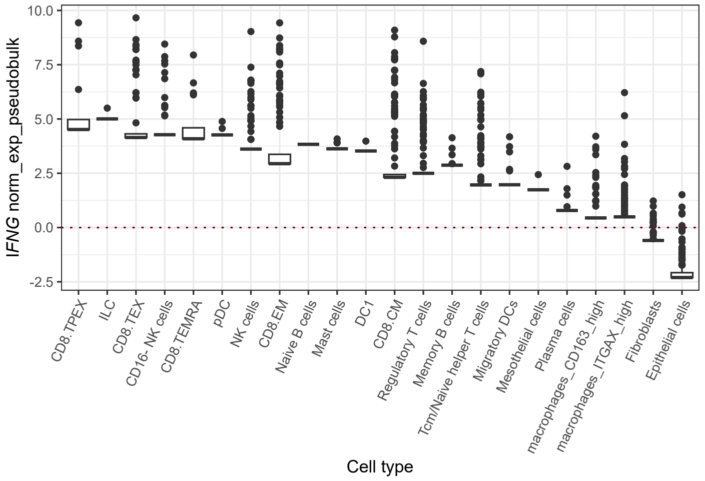
