## Extended Figure 1 for "Single-cell spatial atlas of high-grade serous ovarian cancer unveils MHC class II as a key driver of spatial tumor ecosystems and clinical outcomes"

### Extended material - Figure 1

#### Consensus clustering of cancer cell - ConsensusClusterPlus results

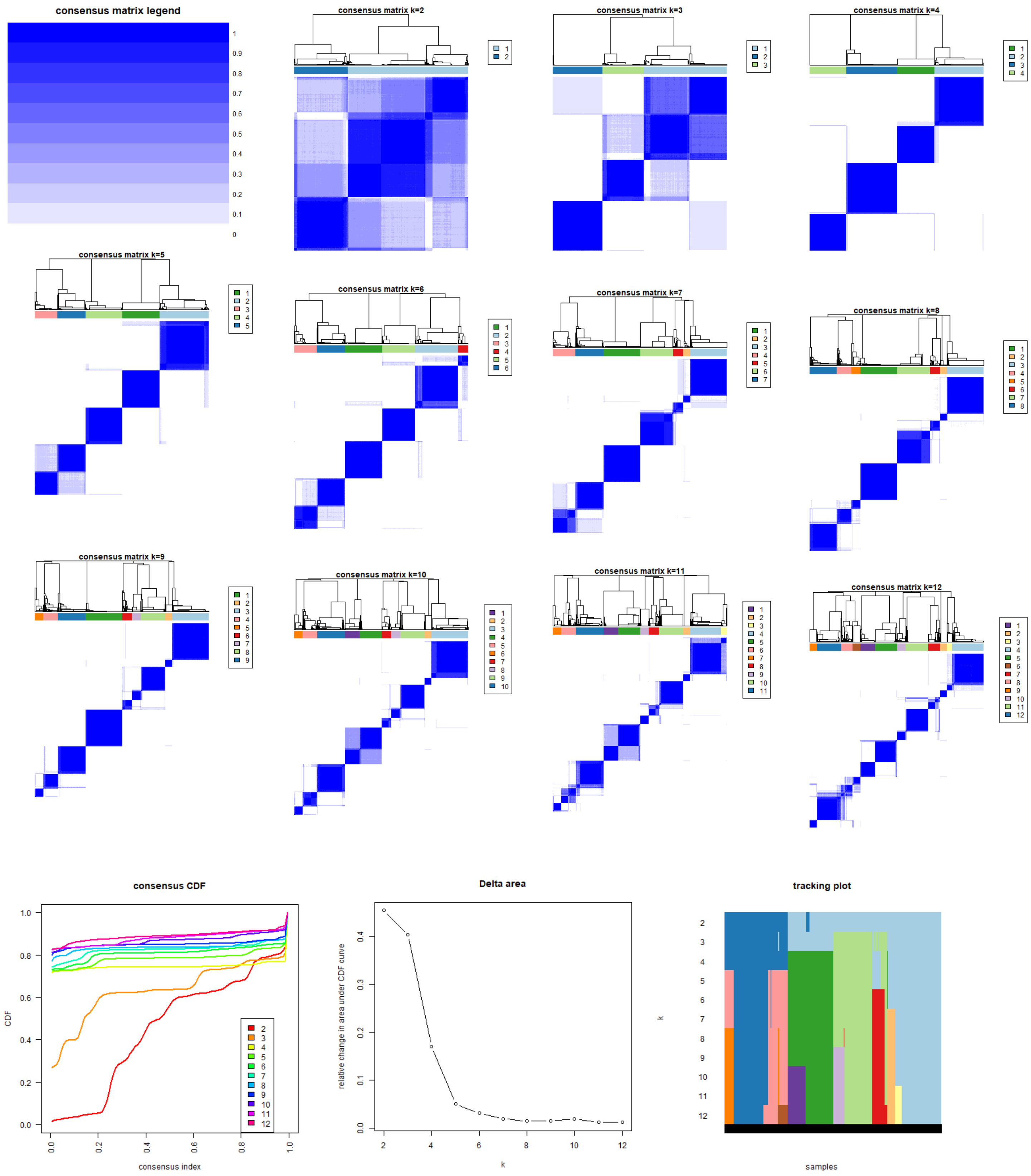
