## Extended Figure 3 for "Single-cell spatial atlas of high-grade serous ovarian cancer unveils MHC class II as a key driver of spatial tumor ecosystems and clinical outcomes"

### Extended material - Figure 3

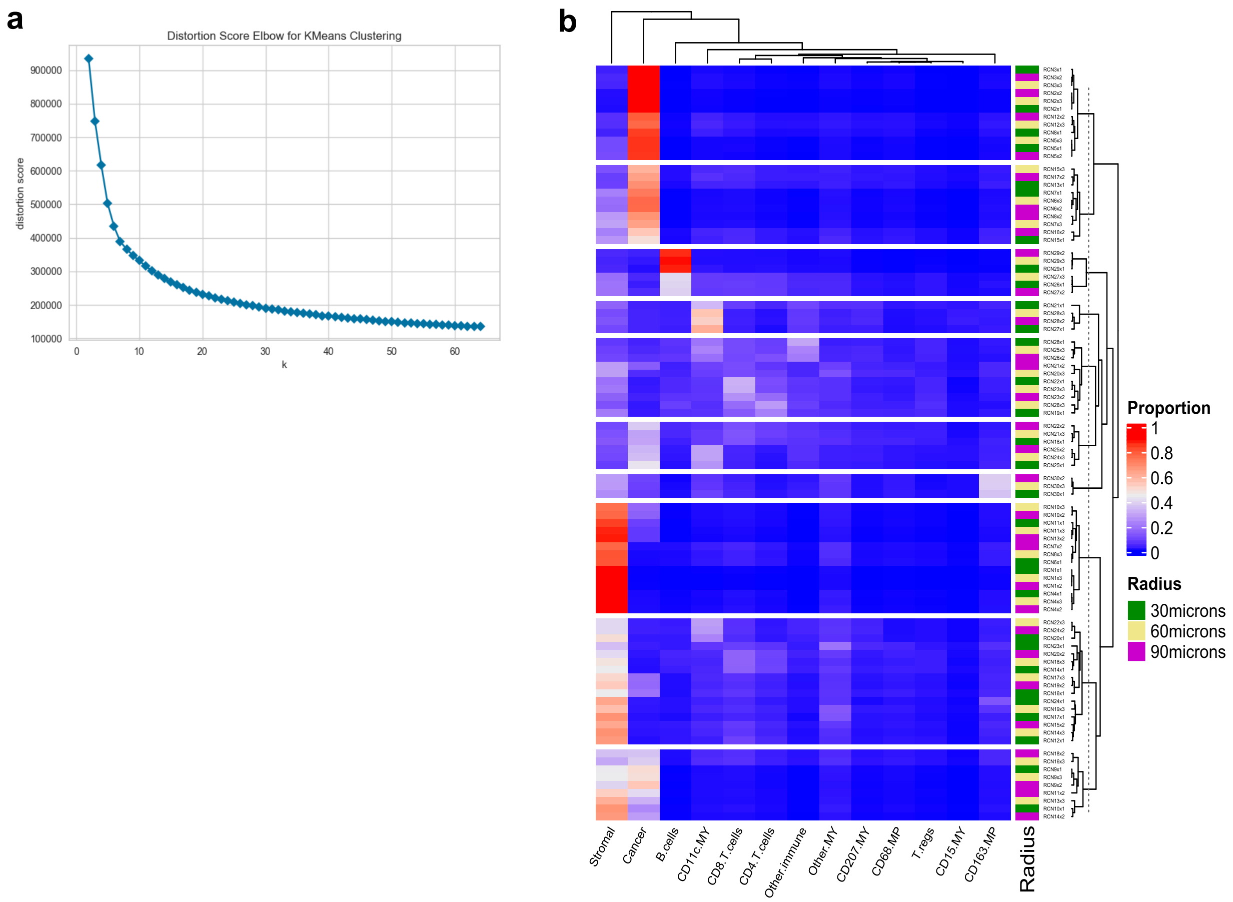


**a** Scatter plot displaying the distortion (sum of squared errors) according to the number of k-means clusters. The cells in the dataset were clustered according to their proportion of neighboring cells in a radius of 30 micrometers. **b** Hierarchical clustering heatmap of the proportion of cell populations per cell neighbor cluster (recurrent cellular neighbors). The cell neighbors were estimated using different radii and then grouped in 30 clusters using k-means. e Heatmap displaying the correlation of RCN abundances across the tumors. Only pairwise significant (p<0.05) correlations are shown.
